# Molecular insights into the capsular polysaccharide transporter Wza-Wzc complex

**DOI:** 10.1101/2025.06.27.662001

**Authors:** Biao Yuan, Christian Sieben, Prateek Raj, Tina Rietschel, Rory Hennell James, Anja Gatzemeier, Lothar Jänsch, Thomas C. Marlovits, Dirk W. Heinz

## Abstract

Capsular polysaccharides (CPS), which form the protective outer capsule surrounding many Gram-negative bacterial pathogens, are critical virulence determinants. Their biosynthesis is primarily carried out *via* the conserved Wzx/Wzy-dependent pathway. In *Escherichia coli*, Group 1 CPS transport through the bacterial envelope is thought to be mediated by the Wza-Wzc complex. In this study, we present the first structural characterization of the complete Wza-Wzc complex from *E. coli* K12, determined using single-particle cryogenic electron microscopy. The structure revealed an elongated, continuous channel spanning the entire envelope, which is crucial for efficient CPS secretion, as supported by mutagenesis studies. Multiple structural snapshots of the ADP-bound Wza-Wzc complex captured intermediate conformations of the double membrane assembly, highlighting its remarkable intrinsic dynamics. In-depth analysis of the isolated Wza translocon and Wzc co-polymerase, revealed new mechanistic details of both complex formation and CPS transport. Importantly, we identified the jellyroll domain of Wzc as a previously unrecognized CPS-binding module, likely guiding CPS repeat units into a proposed Wzc-Wzy polymerization platform. Collectively, these findings provide new structural and functional insights into CPS synthesis and transport, advancing our understanding of bacterial capsule formation and virulence mechanisms.

## INTRODUCTION

The bacterial capsule acts as a thick protective layer around the cell wall, enhancing the bacterium’s ability to withstand environmental stresses. It plays a crucial role in pathogenesis, being involved in virulence, immune evasion, antibiotic resistance, and biofilm formation^1^. Additionally, some of the key components of the capsule, such as capsular polysaccharides (CPS), have significant potential for medical and biotechnological applications^2,3^. CPS synthesis primarily occurs through four assembly pathways involving the Wzx/Wzy-, ABC transporter-, synthase-, or extracellular transglycosylase-dependent mechanisms^4,5^. Of these, the Wzx/Wzy-dependent assembly pathway is the most commonly utilized in both Gram-negative and Gram-positive bacteria, not only for CPS but also for other high molecular weight carbohydrates, such as lipopolysaccharides (LPS), enterobacterial common antigen (ECA), and exopolysaccharides (EPS)^4,6^. *E. coli* CPS K30 and EPS colanic acid are also produced through the Wzx/Wzy-dependent pathway^4,7^. The CPS repeat unit is synthesized in the cytoplasm by a variety of glycosyltransferases, followed by translocation into the periplasm by the flippase Wzx and transfer to the CPS polymerase Wzy for polymerization^6,8–11^.

The CPS co-polymerase Wzc is an inner membrane tyrosine autokinase that regulates CPS chain length during polymerisation^12,13^. Recent cryo-EM structures of *E. coli* K30 Wzc_K540M_, which mimics the dephosphorylated state of the Wzc oligomer, have provided structural insights into its octameric form and domain architecture^12,13^. Specifically, the kinase domain contains the Walker-A ATP-binding motif, which works in conjunction with the tyrosine-rich tail (Y-tail) to facilitate autophosphorylation of these tyrosines^12,13^. The cognate Wzb phosphatase then promotes Wzc oligomerization by dephosphorylating its tyrosine-rich tail, which is localized at the oligomeric interfaces^14,15^.

The jellyroll domain, located between the inner membrane and the periplasmic helical arm domain, has an as-yet-unknown function. In contrast, the periplasmic helical arm domain plays a crucial role for CPS secretion, likely mediating the recruitment of the outer membrane Wza translocon^13^. Orthologous systems in Gram-positive bacteria, such as the CpsC-CpsD protein pair in *Streptococcus pneumoniae*, lack this translocon component, as CPS secretion in these organisms does not require an outer membrane translocon^16,17^.

The coupled secretion machinery is encoded by the *wzabc* operon in *E. coli* K30^7^. Wza can form a stable alpha-helical, “amphora”-like octamer, which is essential for CPS export^18^. The non-phosphorylated Wzc likewise adopts an octameric state, and its interaction with the Wza translocon, validated by *in vivo* cross-linking and negative stain EM analysis, suggests that Wza and Wzc may form a complex for CPS export^13,19,20^.

Despite this, the detailed architecture of the proposed CPS secretion machinery remains unknown. To address this gap, we aimed to provide structural insights into its assembly and the mechanism of CPS secretion. We utilized the orthologous secretion system of colanic acid, which can functionally complement K30 secretion^7^. For simplicity, we refer to colanic acid as CPS hereafter. In this study, we uncovered by cryo-EM that Wzc and Wza form a large secretion machinery across two bacterial membranes. Furthermore, the structure establishes a continuous translocation channel bridging the entire periplasmic space. Multiple conformational states of Wzc, captured both in the presence and absence of Wza, unveil dynamic structural changes involved in the assembly and disassembly of the CPS secretion machinery, marked by significant structural rearrangements of Wzc’s periplasmic helical arms towards the Wza translocon. Additionally, our findings uncover a glycan-recognition role for Wzc’s periplasmic jellyroll domain, providing the molecular basis into the coordinated transfer of CPS repeat units through the Wza-Wzc complex.

## RESULTS

### The Wza-Wzc complex forms a continuous secretion channel

The Wza-Wzc complex has been proposed to be essential for CPS secretion, although with limited *in vivo* evidence^13,20,21^. To address the necessity of an assembled Wza-Wzc complex for CPS secretion *in vivo*, we conducted fluorescence microscopy on live *E. coli* BL21 Star (DE3) cells, overexpressing the entire *wzabc* operon with Wza tagged with mScarlet3 and Wzc tagged with sfGFP. Wza appeared as distinct clusters localized within the membrane, consistent with previous observations for the orthologous Wza from *E. coli* K30 strain^18,20^, suggesting that Wza forms a stable translocon inside the outer membrane. By contrast, Wzc displayed two different populations: distinct clusters and an even membrane distribution, indicating structural heterogeneity and dynamic behavior. The fact that colocalization between Wza (cluster form) and Wzc (cluster form) (**Fig. 1a**) is observed, suggesting a potential complex formation between these two proteins.

**Fig. 1.**
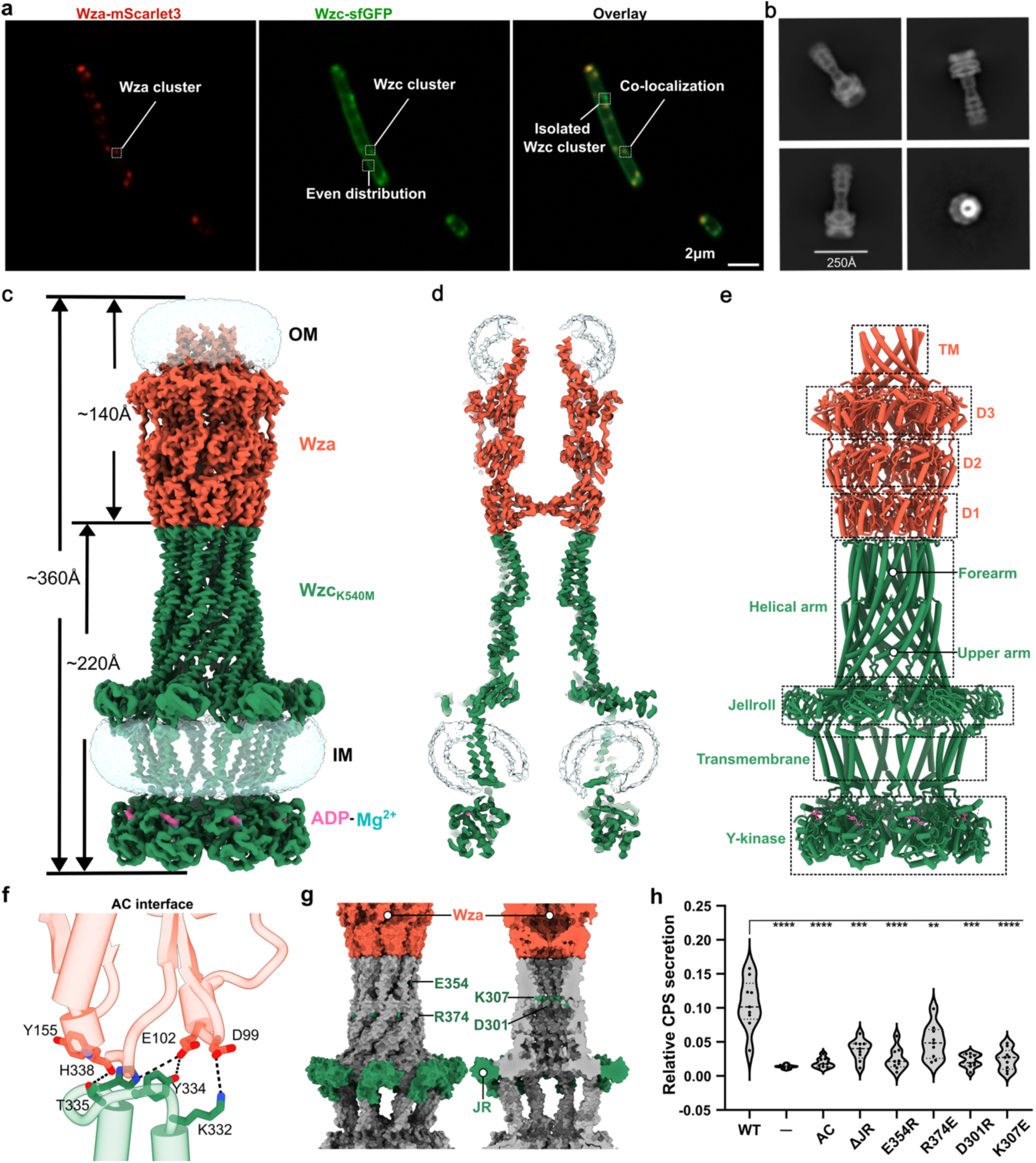
Visualization and structural organization of the Wza-Wzc complex. **(a)** Fluorescence microscopy images of live *E. coli* BL21 Star (DE3) cells expressing the *wzabc* operon from *E. coli* K12. C-terminal fusions of Wza-mScarlet3 and Wzc-sfGFP reveal co-localization between Wza and Wzc. Scale bar: 2 µm. **(b)** 2D class averages of the Wza-Wzc_K540M_ complex. Scale bar: 250 Å **(c)** Cryo-EM density map of Wza-Wzc_K540M_ complex. Bound ADP-Mg^2+^ is labeled. EM density of DDM micelles is shown in transparent blue. **(d)** Cross section of cryo-EM density map of Wza-Wzc_K540M_ complex. **(e)** Atomic model of Wza-Wzc_K540M_ complex with individual subdomains labelled. TM - transmembrane domain; D1, D2, D3 - domains 1, 2, 3; Y-kinase - Tyrosine auto-kinase domain. **(f)** Loop-mediated interactions at the Wza and Wzc_K540M_ interface **(g)** Charged residues located at the oligomerization surface interface and at the narrowest region of the periplasmic channel. **(h)** Mutational analysis of essential structural elements in Wzc. ΔJR indicates deletion of the JR domain (residues 112-204), replaced with a GGGGS linker; AC refers to the AC interface mutant (Wzc_K332E,Y334A,T335A,H338E_). CPS (colanic acid) secretion levels, represented by filled black circles, are averaged from three independent experiments and shown as mean ± s.d.

To further understand the structural architecture of the CPS secretion machinery, we reconstructed the *wzabc* operon under a T7 promoter with the original ribosome binding site sequences and introduced a Strep-tag at the C-terminus of Wza. Only a small amount of Wzc and Wzb co-purified with Wza (**Fig. S1a**), consistent with our live-cell imaging data showing a substantial portion of Wzc was evenly distributed in the inner membrane and did not colocalize with Wza (**Fig. 1a**). By introducing the K540M mutation to *E. coli* K12 Wzc, we were able to purify a stable *E. coli* K12 Wza-Wzc_K540M_ complex (**Fig. S1b, c**). K540 is a central residue within the conserved Walker-A motif responsible for ATP binding and hydrolysis^22,23^. Mutating it to methionine leads to stable Wzc octamer formation in *E. coli*^13^.

Cryo-EM analysis of the Wza-Wzc_K540M_ complex, with a global spatial resolution of 3.5 Å (without symmetry applied) and 3.2 Å (applying C8 symmetry), revealed a well-defined, channel-shaped assembly spanning both the inner and outer membranes (**Fig. 1b, c and Fig. S2**). These reconstructions demonstrated the highly symmetric nature of the complex with a length of approximately 360 Å, composed of the outer membrane Wza translocon (140 Å) and the inner membrane tyrosine auto-kinase Wzc_K540M_ (220 Å). This state is designated as the Conformation 0 (Conf 0), representing the ADP-Mg^2+^ bound Wza-Wzc complex.

Segmented representation showed a central cavity, characteristic of the CPS secretion machinery (**Fig. 1d**). In the periplasmic region (**Fig. 1e**), the complex comprised Wza domains D1-D3 and Wzc’s jellyroll (JR) domain and helical arm (HA) domain consisting of an upper arm and a forearm. The extended HA forearm interacts with Wza D1 to form a continuous CPS transport path. The ADP-Mg^2+^ bound Y-kinase domain forms the cytoplasmic region of the complex, and mediates its disassembly via auto-phosorylation^15^.

To investigate how the structural features of Wzc influence CPS secretion, we conducted site-directed mutagenesis on wild type Wzc targeting four critical regions: the Wza-Wzc (AC) interface (D99-K332, E102-Y334, E102-H338, Y155-T335), the JR domain, charged surface residues at the oligomeric interfaces (E354, R374), and charged channel residues within the narrowest part of the Wzc channel (D301, K307) (**Fig. 1f, g**). To assess the functional impact of the mutations on CPS secretion, we utilized a standard colanic acid secretion assay using an *E. coli* JM109 (DE3) Δ*wzabc* strain complemented with plasmids expressing the *wzabc* operon carrying the respective Wzc variants^24^. Quantitative analysis revealed a significant decrease in CPS secretion for all mutants (**Fig. 1h**). Notably, the AC interface mutations (Wzc_K332E,Y334A,T335A,H338E_) completely abolished CPS secretion, underscoring the essential role of the Wza-Wzc complex in the CPS secretion process. Deletion of the JR domain led to marked but incomplete loss of secretion. Additionally, charge-reversal mutations at the oligomeric interface (E354R, R374E) and within the inner channel (D301R, K307E) impaired CPS secretion, suggesting that a proper channel architecture within the HA domain is required for efficient CPS transport.

### Cryo-EM structures reveal different conformations of CPS secretion machinery

Magnesium ions (Mg^2+^), essential cofactors for ATP hydrolysis, play a critical role in stabilizing both the enzyme-ADP complex and the transition state during the reaction^15,25,26^. Additionally, Mg^2+^ facilitates oligomerization of the Wzc kinase domain^25^. Interestingly, ADP alone binds to the Wzc kinase domain with approximately sixfold greater affinity than the ADP-Mg^2+^ complex^25,26^. As a result, chelation of Mg^2+^ by EDTA could likely capture distinct transient states of the ADP-bound Wza-Wzc complex. When 10 mM EDTA was added to remove Mg^2+^, 2D classification in fact revealed a significant increase in particles consisting of a single Wzc octamer associated with two Wza translocons compared to the untreated sample (**Figs. S2b and S3b**). Subsequent 3D reconstructions identified five additional conformations (Confs I − V) of the Wza-Wzc complex (**Fig. 2 and Fig. S3**), highlighting the dynamic nature of the CPS secretion machinery during the secretion process.

**Fig. 2.**
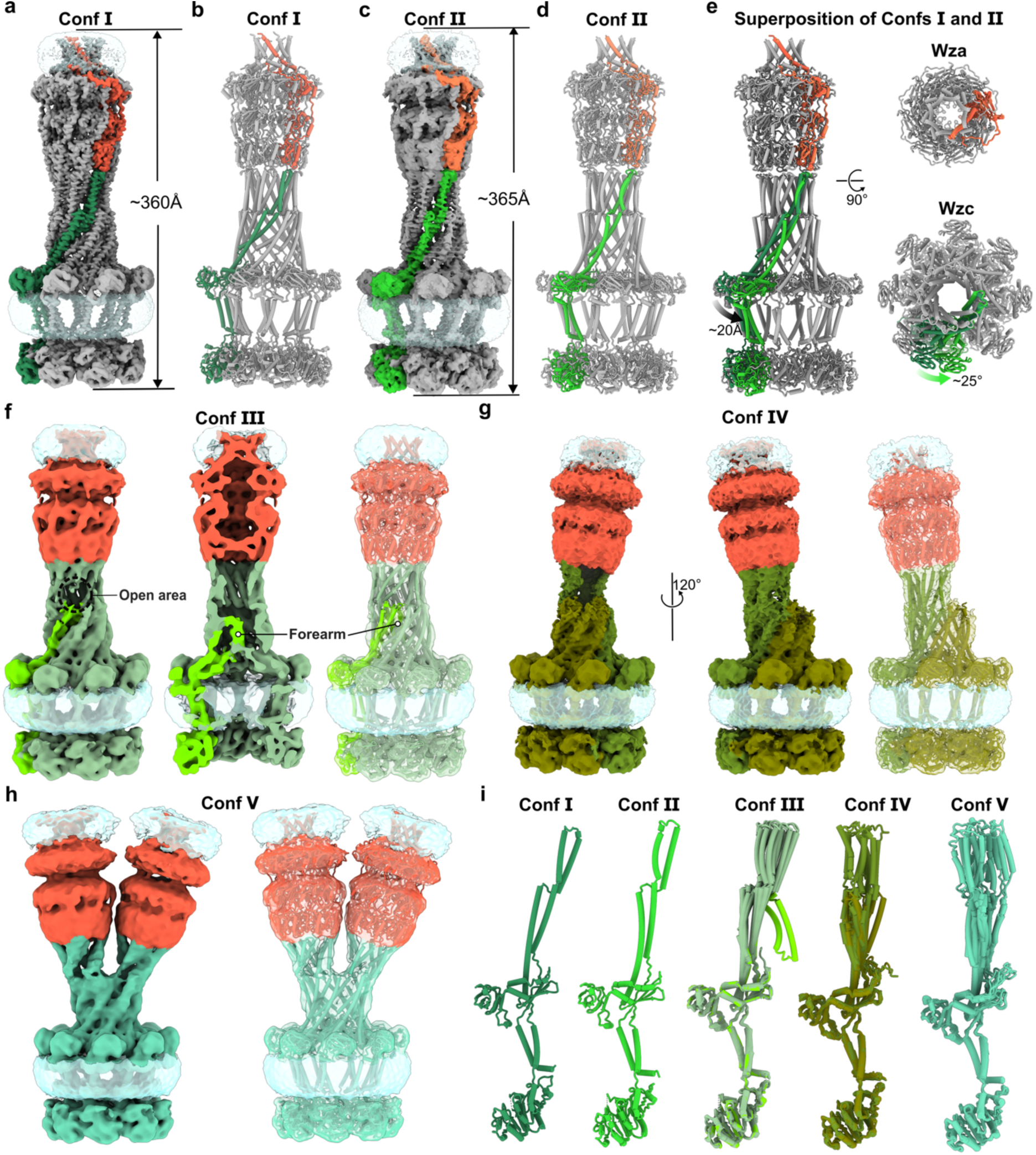
Dynamics of the Wza-Wzc complex. **(a)** Cryo-EM density map of Conf I Wza-Wzc_K540M_ complex. The density corresponding to a single Wza protomer is colored in tomato red, while the interacting Wzc protomer is shown in sea green. “Conf” denotes conformation. **(b)** Structural model of Conf I Wza-Wzc_K540M_ complex. **(c)** Cryo-EM density map of Conf II Wza-Wzc_K540M_ complex. The EM densities of a Wza protomer and its interacting Wzc protomer are shown in coral and lime green, respectively. **(d)** Structural model of Conf II Wza-Wzc_K540M_ complex. **(e)** Structural superposition of Conf I and Conf II the Wza-Wzc_K540M_ complexes, aligned on the Wza translocon to highlight conformational differences. **(f)** Cryo-EM density map of Conf III Wza-Wzc_K540M_ complex showing a partially open periplasmic channel with a cross-section revealing one HA forearm protruding into the channel interior. **(g)** Cryo-EM density map of Conf IV Wza-Wzc_K540M_ complex, characterized by a large opening at the interface between Wzc and Wza. **(h)** Cryo-EM density map of Conf V Wza-Wzc_K540M_ complex, where a single Wzc octamer is associated with two Wza translocons. **(i)** Comparison of Wzc protomers across Confs I − V. Conf I shows the protomer from the Conf I complex; Conf II shows the protomer from the Conf II complex; Conf III − V shows a superposition of all protomers from each respective state.

The global resolution of the Conf I structure decreased to 3.8 Å compared to that of Conf 0 (C1, **Fig. S3**). Structurally, Conf I closely resembled the Conf 0 shown in **Fig. 1c** and, as anticipated, lacked Mg^2+^ in the ADP-binding pocket (**Fig. 2a, b and Fig. S4a**). This absence appeared to trigger a subtle cascade-like motion propagating from the Y-Kinase domain to the HA domain (**Fig. S4b**). Conf II revealed an expanded CPS secretion machinery, approximately 5 Å longer than Conf I (**Fig. 2c, d**). This is primarily driven by twisting of the HA domain, resulting in a ∼25° rotation of Wzc (**Fig. 2e**). This elongation likely generates a mechanical force sufficient to extract the final CPS repeat unit from the inner membrane, thereby aiding its dissociation from Wzy. Conf III exhibited a partially open periplasmic channel, with one HA forearm positioned inside the incompletely sealed channel (**Fig. 2f**). In Conf IV, the periplasmic channel was largely open, and the Wza translocon appeared almost detached from the Wzc octamer (**Fig. 2g**). In this conformation, the Wza translocon remained associated with Wzc via just four HA domains, suggesting that four Wzc protomers are sufficient to maintain Wza association. The remaining four forearms, which were not interacting with Wza, likely became more flexible, and were therefore unresolved in the cryo-EM map. Finally, Conf V revealed a configuration in which the HA domains split into two halves (opening angle ∼60°), with four protomers in each half associated with a separate Wza translocon (**Fig. 2h**). This finding further confirmed that four accessible HA domains are sufficient to recruit and engage a Wza translocon. Collectively, these results demonstrate the intrinsic flexibility of the Wzc HA domains, particularly in the forearm regions (**Fig. 2i**), which plays a crucial role in the assembly and disassembly process of CPS secretion machinery.

### Novel insights into the assembly of the Wza-Wzc complex

The Wza-Wzc complex forms an octameric trans-envelope structure essential for CPS secretion in most Gram-negative bacteria. This large, dynamic assembly spans both membranes, with the observed conformational flexibility of the Wzc HA domain playing a pivotal role in coordinating CPS polymerization and export through the periplasm. To elucidate how Wza and Wzc initiate assembly of this nanomachine, we determined the cryo-EM structures of the isolated Wza translocon at a resolution of 3.0 Å (C1) and 2.7 Å (C8) (**Fig. S5**), along with the structures of the isolated Wzc_K540M_ octamer at 3.4 Å (C1) and 3.2 Å (C4) resolution (**Fig. S6**).

The Wza translocon maintained the same conformation in its isolated octameric form as it did when complexed with Wzc (**Fig. 3a-c**), indicating high structural rigidity and stability across different functional states of the complex. In contrast, the isolated Wzc_K540M_ octamer exhibited marked conformational diversity, existing in two distinct protomeric states within the octamer. In Conformation i (Conf i), four protomers assembled into a V-shaped channel, with the forearms of the HA domains tightly aligned. In Conformation ii (Conf ii) the upper arms of four protomers rotated outward, leaving the forearms more flexible and unresolved in the structure (**Fig. 3d-g**). This suggests that in Conf ii, the forearms become exposed and dynamic, unlike their shielded position in Conf i (**Fig. 3f, g**), allowing Wzc to engage and recruit the Wza translocon.

**Fig. 3.**
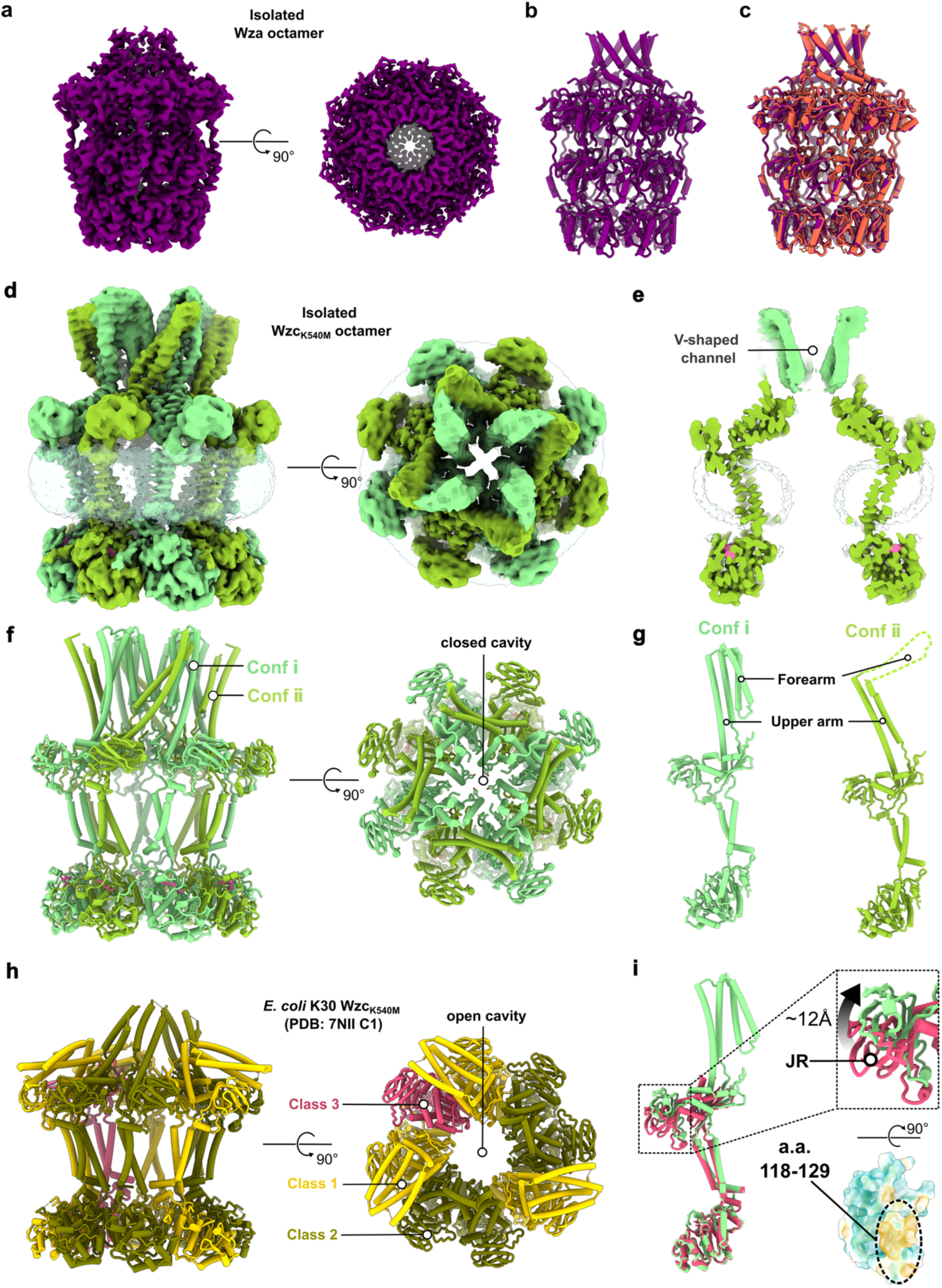
Cryo-EM structures of the isolated Wza translocon and Wzc octamer. **(a)** Cryo-EM density map of the Wza translocon resolved with C8 symmetry. **(b)** Structural model of the Wza translocon. **(c)** Structural comparison of Wza before (purple) and after (tomato red) complex formation with Wzc, illustrating its conformational stability. **(d)** Cryo-EM map of the isolated *E. coli* K12 Wzc_K504M_ octamer under C4 symmetry. **(e)** Cross-sectional view showing the inner membrane cavity of *E. coli* K12 Wzc_K504M_ sealed by the HA domains in Conf i of Wzc_K504M_. **(f-g)** Structural models of *E. coli* K12 Wzc_K504M_ octamer. Conf i protomers are colored light green; Conf ii protomers are yellow green. Only the HA forearm of the Conf ii protomer is resolved. (**h**) Model of *E. coli* K30 Wzc_K540M_ (C1 symmetry; PDB: 7NII)^13^. (**i**) Superimposition of *E. coli* K12 Conf i Wzc_K540M_ and *E. coli* K30 Wzc_K540M_ (class 3) protomers reveals vertical displacements of the JR domain. The hydrophobic helix (residues 118-129) within the JR domain lies near the inner membrane in the class 3 conformation, suggesting membrane association. Upward movement of the JR domain during conformational transitions (class 3 to Conf i) likely facilitates its dissociation from the membrane, potentially contributing to the CPS translocation.

For complete translocation channel formation with Wza, Wzc must undergo a rotational conformational shift toward the outer membrane, extending all forearms to establish a sealed translocation pathway (**Fig. S7a, b**). Notably, the periplasmic domain of Wzc (Wzc_Peri_), comprising the HA- and JR-domains, alone was unable to form a complex with Wza (**Fig. S7c**), supporting a model in which oligomerization of the cytoplasmic Y-kinase domain drives assembly by exposing HA forearms necessary for Wza engagement. This mechanism supports previous findings^13,19,22^ linking Y-kinase domain-driven transitions to Wza-Wzc complex formation, regulated by the phosphorylation state of the Y-tail. To dissect this relationship, we analysed the phosphorylation state of the recombinantly produced wild-type Wzc by mass spectrometry. Wild-type Wzc existed as a lower-order oligomeric form *in vitro* (**Fig. S7d**). Mass spectrometry analysis demonstrated that while up to four tyrosines can undergo simultaneous phosphorylation, with the peptides bearing one (1pY) or two (2pY) phosphorylated tyrosines predominated (**Fig. S7e**). The single phosphorylation Y705 was found most abundant followed by Y708 and Y715. Association analyses indicated higher order phosphorylations have prevalence to involve the sites Y710, Y713 and Y715 (**Fig. S7f)**. In total, 43 different phosphorylation site combinations (with phosphorylation probabilities above 0.85) were detected, suggesting a redundant phosphorylation pattern along dynamic assembly (**Table S1**). This redundancy aligns with the observation that no single phosphorylation site is indispensable for CPS secretion^27^.

Compared to the previously reported *E. coli* K30 Wzc_K540M_ structure^13,28^, the *E. coli* K12 Wzc_K540M_ exhibited a preference for C4 symmetry, with its inner membrane cavity sealed by the forearms in Conf i (**Fig. 3h and Fig. S7g, h**). This closed arrangement likely represents the native *in vivo* state, serving to prevent ion leakage across the membrane before forming the complex with Wza. Superimpositions of the Wzc protomers from the *E. coli* K30 and K12 strains indicated that the JR domain might move vertically relative to the inner membrane (**Fig. 3i**). Additionally, the presence of a hydrophobic helix (residues 118-129) within the JR domain (**Fig. 3i**), located near the inner membrane, suggests transient lateral membrane association that may be disrupted during conformational transitions.

### Wzc JR domain enables CPS repeat unit recognition and loading

Although we resolved the Wza-Wzc complex required for CPS polymerization and export, how CPS is loaded into this machinery and in particular, how Wzc and Wzy interact during CPS polymerization remain unclear. Recent structural and biochemical studies of the homologous WzzE-WzyE (co-polymerase-polymerase) complex, involved in polymerization of the enterobacterial common antigen, suggest the formation of a polysaccharide polymerization platform with an 8:1 stoichiometry between co-polymerase and polymerase^10^. Weckener et al. hypothesized that this structural arrangement might also apply to its homologous CPS polymerization system where Wzc functions as the co-polymerase^10^. To further test this hypothesis, we used AlphaFold3 ^29^ to predict a putative Wzc-Wzy (ratio 8:1) complex structure within the inner membrane. The prediction suggests a similar inner membrane platform for CPS polymerization (interface predicted Template Modelling (ipTM) score: 0.56) (**Fig. 4a**).

**Fig. 4.**
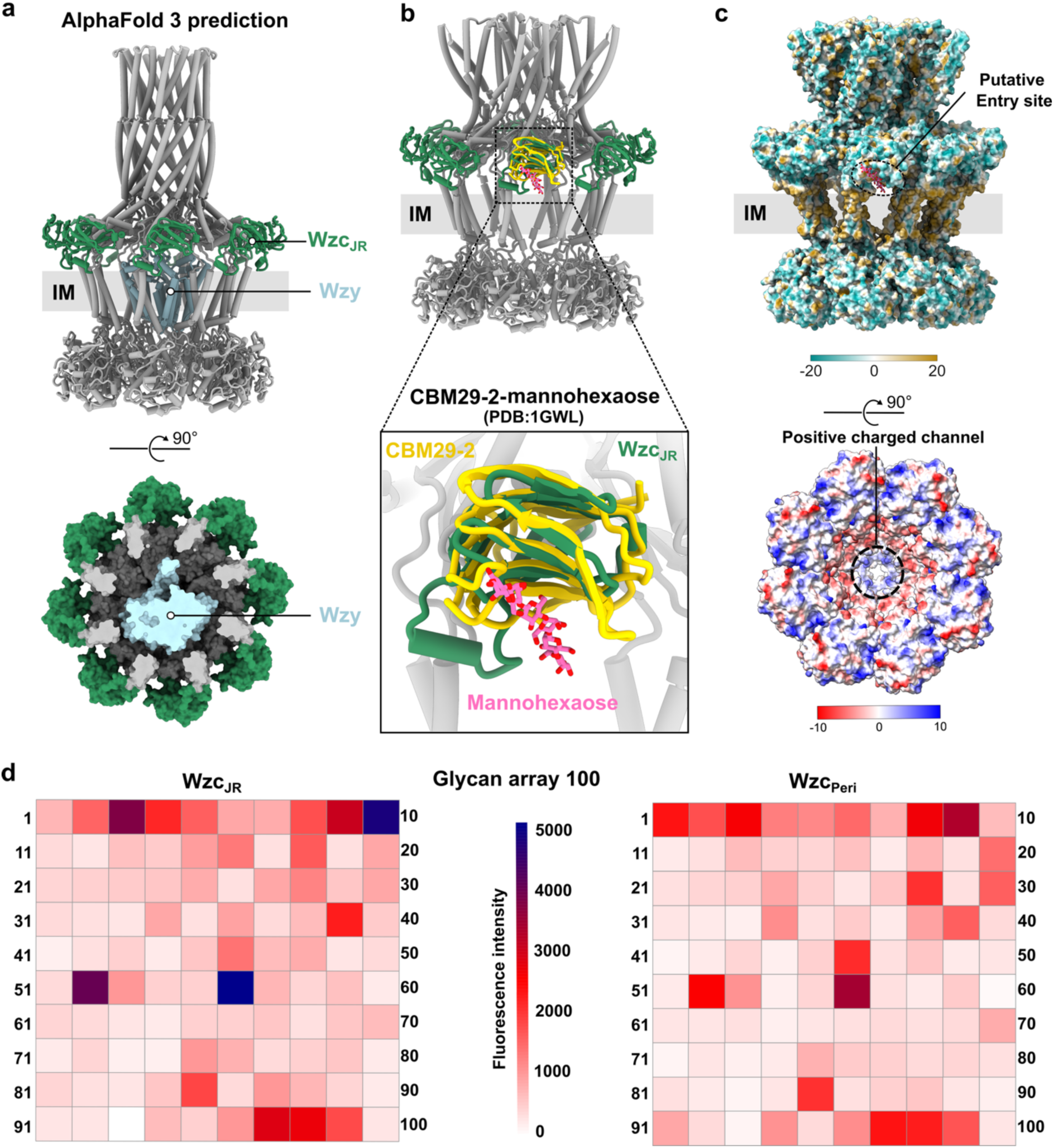
The CPS polymerization platform Wzc–Wzy with JR-directed CPS loading. **(a)** AlphaFold3 model of the Wzc-Wzy complex. Side (top panel) and bottom (bottom panel) views of the predicted structure reveal that Wzc inner membrane cavity is sufficiently spacious to accommodate a single Wzy molecule. The co-polymerase Wzc JR domain (Wzc_JR_) is colored in sea green, and the polymerase Wzy is colored in light blue. **(b)** Superimposition of the Wzc_K540M_ structure with the mannohexaose bound structure of CBM29-2 (PDB: 1GWL)^31^. **(c)** Surface properties of the Wzc_K540M_ structure reveal a putative CPS repeat unit entry site and a positively charged CPS polymerization channel. Surface hydrophobicity highlights the proximity of the superimposed mannohexaose to the putative CPS repeat unit entry site, formed by JR, TM, and inner membrane plane (Top panel). Surface electrostatic potential of the Wzc_K540M_ octamer reveals a positively charged channel likely guiding CPS polymerization towards the Wza translocon. The transmembrane region is labelled as IM. JR is labelled as Wzc_JR_. **(d)** Glycan binding profiles of *E. coli* K12 Wzc_JR_ and Wzc_Peri_ (Wzc without TM and Y-kinase domains) using Glycan Array 100 (Raybiotech). Cys3-conjugated Streptavidin were used to detect the binding ability of biotinylated Wzc_JR_ and Wzc_Peri_. The relative fluorescence intensity of Wzc_JR_ (25 µg/ml) and Wzc_Peri_ (50 µg/ml) binding to each glycan printed on the glass slide provided from Glycan Array 100 kit is presented in a heatmap format. The scale bar is from 0 to 5000.

This positioning of Wzy within the inner membrane cavity of Wzc (**Fig. 4a**) raises the question of how CPS repeat units are subsequently delivered to the polymerization platform after their translocation across the inner membrane by the flippase Wzx. Strikingly, the JR domain of Wzc, with a yet unknown function, is situated near the periplasmic side of the inner membrane. The jellyroll fold is a structural motif commonly found in various carbohydrate binding proteins including L-type lectins, carbohydrate binding modules (CBMs), and members of the glycoside hydrolase family 16 (GH16)^30–32^. Positioned near both the membrane and the core of the proposed Wzc-Wzy polymerization platform, this domain is well-placed to facilitate the transfer of membrane-anchored CPS repeat units from the flippase Wzx to the polymerase Wzy. Importantly, deletion of the JR-domain nearly abolished CPS secretion (**Fig. 1**), indicating its critical role in coupling CPS to the secretion machinery.

Based on these structural analyses, we propose that the JR domain acts as a recognition module for CPS repeat units, aiding in their transfer from Wzx to the CPS polymerization platform. Superposition of the JR domain of Wzc_K540M_ with the crystal structure of CBM29-2 bound to hexasaccharide mannohexaose^31^ shows a potential CPS repeat unit binding site in the JR domain. The location of the mannohexaose in this superposition suggests an analogously bound membrane anchored CPS repeat unit interacting in a similar fashion with the JR domain of Wzc (**Fig. 4b**). The positioning of the CPS repeat unit within the Wzc octamer further reveals a potential substrate entry site towards the Wzc-Wzy polymerization platform. This site is formed by the inner membrane plane along with the TM and JR domains of two neighboring Wzc protomers and appears to be sufficiently large (∼15 Å high and ∼24 Å wide) for CPS repeat units entry (**Fig. 4c**). We therefore propose that CPS repeat units enter the polymerization platform through this site, where the JR domain captures and directs them into the inner membrane cavity for polymerization by Wzy. Polymer growth then proceeds toward the Wza translocon, guided by the positively charged HA channel of Wzc (**Fig. 4c**).

To further assess whether the JR domain is capable of recognizing CPS, we conducted a glycan array screen using both the isolated JR domain and Wzc_Peri_ at three different protein concentrations (**Fig. 4d and Fig. S8**). Notably, both constructs bound a similar range of glycans, identifying the JR domain as the minimal structural element responsible for CPS recognition.

## DISCUSSION

Synthesis and transport of CPS *via* the Wzy-Wzx-dependent pathway plays a fundamental role in the virulence, environmental adaptability, and immune evasion strategies of most Gram-positive and Gram-negative bacteria^33^. In this study, we present the first complete structure of the CPS secretion machinery formed by the *E. coli* K12 Wza-Wzc complex. The structure, which is an octameric heterodimer, reveals a 360 Å long trans-envelope channel that spans the inner and outer membranes. Multiple structural snapshots of the Wza-Wzc complex, along with the isolated Wza and Wzc octamers, offer important insights into the assembly and disassembly dynamics of the CPS secretion machinery.

While the Wza translocon maintains a strikingly stable conformation during these transitions, the Wzc co-polymerase exhibits substantial conformational rearrangements, particularly within its helical arm (HA) domain. This domain transitions from a rather compact form to an extended state, allowing it to reach and interact with Wza for complex formation. Resolved structures of the Wza-Wzc complex also reveal distinct conformational states, emphasizing the intrinsic flexibility and dynamic coordination required for CPS transport. The HA domain’s structural variability highlights its central role in modulating CPS polymerization and export. Notably, in the absence of Mg^2+^, conformational changes are initiated, propagating from the cytoplasmic Y-kinase domain to the HA domain. This results in a dramatic transition from a stable, continuous secretion channel to a widely open conformation, revealing a potential mechanical basis for CPS extrusion and subsequent disassembly of the CPS secretion machinery. A particularly interesting finding is the observation of a Wzc octamer associated with two Wza translocons, suggesting the potential for alternative assembly states. This dual-translocon configuration may represent a mechanism for enhancing CPS secretion efficiency by dynamically switching the HA domain interaction between two translocons. Further studies, including *in vivo* validation, will be essential to determine whether such configurations occur naturally, thus contributing to CPS export in bacterial systems.

While our cryo-EM and biochemical data offer unprecedented insight into the Wzx/Wzy-dependent synthesis pathway, the broader evolutionary context and mechanistic distinctions among bacterial polysaccharide transport pathways merit further consideration. In the ABC-transporter-dependent pathway, CPS is synthesized within the cytoplasm and translocated across the inner membrane by the ABC transporter complex^4^. A recent cryo-EM structure of KpsMT bound to its periplasmic adaptor KpsE revealed a potential secretion pathway through the inner membrane^34^. Moreover, AlphaFold modeling of the KpsMT-KpsE in complex with outer membrane translocon KpsD bears striking structural resemblance to the Wza-Wzc machinery characterized in our study^34^. This convergence implies a common evolutionary mechanism across Gram-negative species to develop large, contiguous secretion supercomplexes that enable the efficient and tightly regulated export of CPS. Further comparative and phylogenetic analyses will be instrumental in unraveling whether these systems arose via convergent or divergent evolutionary trajectories.

Unlike the relatively simple O-antigens, CPSs often display large chemical and structural diversity, frequently incorporating a broad repertoire of monosaccharides, amino sugars, and occasionally non-carbohydrate moieties such as pyruvate or acetyl groups^1,35^. This structural diversity enables bacteria to tailor their capsules to diverse ecological niches and host immune environments^1,35^. Intriguingly, this complexity is mirrored in the architecture of their co-polymerase. The O-antigen co-polymerase Wzz lacks JR and Y-kinase domains^13,28^ (**Fig. S9**). In contrast, the JR domain of CPS co-polymerase Wzc can bind a wide range of glycans, providing the molecular basis for CPS diversity and specificity. The functional flexibility of the JR domain likely underlies the ability of the Wza-Wzc system to accommodate various CPS types, as exemplified by its capacity to mediate the export of both colanic acid and K30 CPS via *E. coli* K12 Wza-Wzc complex^7^.

Our data further support a phosphorylation-dependent mechanism that regulates Wzc oligomerization^14,19,22^. Phosphorylation analysis of wild type Wzc indicates that the Y-tail undergoes multiple phosphorylation events, with Y710, Y713, and Y715 being the most frequently modified among higher order phosphorylations. These residues are situated at the oligomeric interface of the Wzc octamer, supporting a model in which autophosphorylation regulates its dynamic assembly. These findings broaden our understanding on the involvement of transient phosphorylation pattern during the dynamic assembly of Wzc. Dephosphorylation of the C-terminal Y-tail triggers the Y-kinase domain octamerization^14,19,22^, which further promotes the rearrangement of HA domains to engage the Wza translocon. Wzz, lacking the Y-kinase domain, instead forms an octameric assembly via its HA domain^22,36^ (**Fig. S9**). Although the HA domain architecture is broadly conserved between Wzc and Wzz, Wzc’s HA is significantly extended^28^, suggesting functional divergence. This elongation, along with the presence of the additional regulatory domains, underscores the specialized adaptations of Wzc to support CPS assembly and export via the Wza translocon, distinguishing it from Wzz’s role in lipopolysaccharide assembly and secretion.

Our findings support a comprehensive model for CPS polymerization and export (**Fig. 5**). Following the SecYEG-SPII maturation of Wza^37^, the protein assembles into a stable octameric translocon in the outer membrane, independently of other components^18^. In contrast, Wzc predominantly exists in a monomeric or heterogeneous oligomeric state when phosphorylated. Dephosphorylation of the tyrosine residues in the C-terminal Y-tail by the cognate phosphatase Wzb triggers the assembly of Wzc into a stable octamer likely surrounding the inner membrane polymerase Wzy^10,33^. Conf i HA domains form a polymerization platform with Wzy, directing CPS chain elongation toward the Wza translocon. Meanwhile, the flexible Conf ii HA domains facilitate recruitment of the Wza translocon. Positioned at the periplasmic face of the inner membrane, the JR domain acts as a glycan receptor, capturing CPS repeat units flipped by Wzx and delivering them to Wzy for polymerization. The helical movement generates a mechanical force through HA domain twisting that drives extrusion of the CPS polymer from the inner membrane to the Wza translocon. Upon completion of translocation, the Wzc octamer dissociates through autophosphorylation, effectively resetting the machinery for a new cycle of CPS biosynthesis and export.

**Fig. 5.**
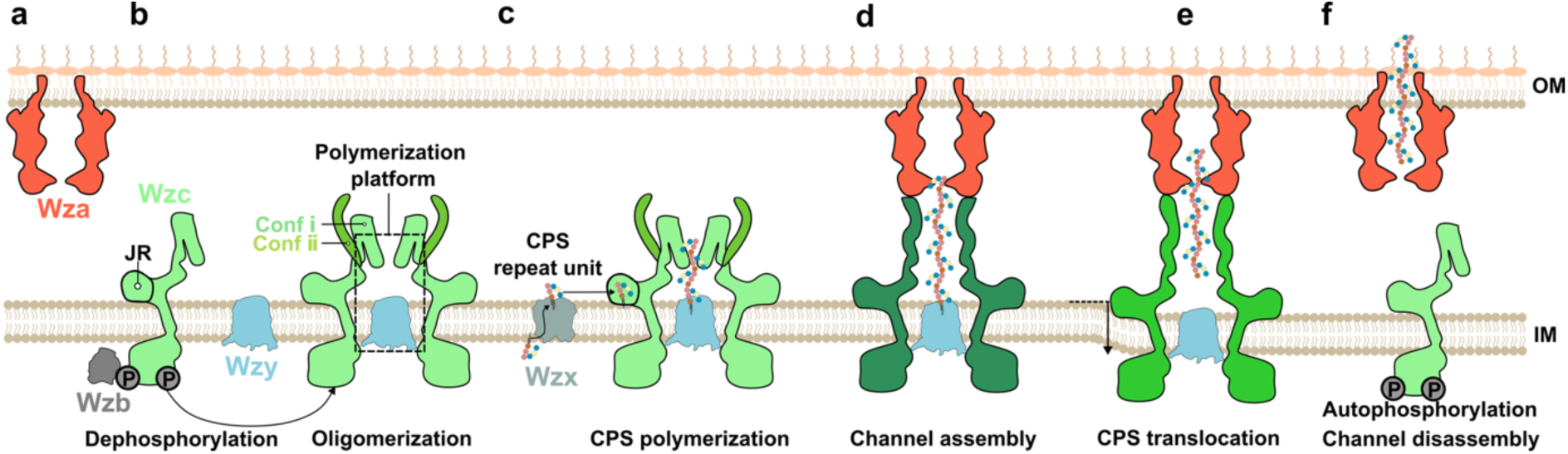
Proposed model of CPS polymerization and secretion. **(a)** The Wza translocon is pre-assembled as an octamer on the OM**. (b)** In its phosphorylated state, Wzc remains monomeric, with the HA domain locked in a conformation that prevents recruitment of the Wza translocon. Dephosphorylation by the Wzb phosphatase induces Wzc oligomerization, likely around the CPS polymerase Wzy, leading to the formation of CPS polymerization platform. **(c)** CPS repeat units, translocated from the cytoplasm to the periplasm by the flippase Wzx, are recognised by the JR domain and subsequently delivered to Wzy within the transmembrane cavity of Wzc for polymerization. **(d)** Once the CPS chain reaches maturity, the HA domains open, forming a continuous secretion channel with Wza translocon that spans the entire periplasm. **(e)** The twisting motion of the HA domains, which extends the secretion channel, may generate the mechanical force necessary to release the CPS chain from Wzy into the Wza translocon. **(f)** The mature CPS is then exported across the outer membrane *via* the Wza translocon, followed by disassembly of the secretion machinery through Wzc auto-phosphorylation.

Together, our structural and biochemical data reveal a highly dynamic CPS secretion machinery, in which coordinated domain rearrangements and glycan recognition converge to ensure efficient capsule biosynthesis and export. This work advances our understanding of bacterial virulence and may inform the development of novel antimicrobial strategies targeting polysaccharide transport systems.

## ACKNOWLEDGMENTS

We thank the Heinz group members: Surabhi Lata and Ines Glökner for support of this project. We thank Sven-Kevin Hotop and Michael Lehky for supporting the scanning glass slides from the glycan array. We also thank Dominik Körner and the Core Facility for Translational Proteomics (CFTP, HZI) for generating and providing the phosphoproteome data and Frank Klawonn for statistical evaluation of phosphosite analyses. Furthermore, we thank Stefan Schmelz for maintaining the cryo-EM facility. We thank Konrad Büssow, Youssef El Mouali Benomar, and Joop Van den Heuvel for kindly providing some plasmids. We thank Nathan Nagar for help with large AlphaFold 3 predictions. This project was supported in part through funds available to T.C.M. through the Behörde für Wissenschaft, Forschung und Gleichstellung of the city of Hamburg at the Institute of Microbial and Molecular Sciences at the University Medical Center Hamburg-Eppendorf (UKE) and the Deutsches Elektronen Synchrotron (DESY). C.S. acknowledges support by the Helmholtz Association (VH-NG-1526).

## AUTHOR CONTRIBUTIONS

Conceptualization: B.Y., T.C.M., D.W.H., cryo-EM sample preparation, data-collection, processing, and model building: B.Y., Structural analysis: B.Y., D.W.H., R.H.J., T.C.M., Fluorescence microscopy: C.S., B.Y.. Mass spectrometry: T.R., B.Y., L.J.. Glycan array: B.Y., P.R.. Cloning and CPS secretion assay: B.Y., A.G.. Visualization: B.Y., D.W.H., D.K., C.S., R.H.J., L.J., T.C.M.. The original draft: B.Y., D.W.H.. Reviewing and editing: all authors.

## MATERIAL AND DATA AVAILABILITY

Models and associated cryo-EM maps have been deposited into the PDB database with the following accession codes: Wza_C1 (EMD-53598; PDB: 9R60), Wza_C8 (EMD-53599; PDB: 9R61), Wzc_K540M__C1 (EMD-53600; PDB: 9R62), Wzc_K540M__C4 (EMD-53601; PDB: 9R63), Wza-Wzc_K540M__C1 (Conf 0; EMD-53602, PDB: 9R64), Wza-Wzc_K540M__C8 (Conf 0, EMD-53603, PDB: 9R65), Wza-Wzc_K540M__C1 (Conf I; EMD-53604, PDB: 9R66), Wza-Wzc_K540M__C8 (Conf I; EMD-53605; PDB 9R67), Wza-Wzc_K540M__C1 (Conf II; EMD-53606; PDB 9R68), Wza-Wzc_K540M__C8 (Conf II, EMD-53607; PDB 9R69), Wza-Wzc_K540M__C1 (Conf III; EMD-53608; PDB 9R6A), Wza-Wzc_K540M__C1 (Conf IV; EMD-53609; PDB 9R6B), Wza-Wzc_K540M__C1 (Conf V; EMD-53610; PDB 9R6C). All the raw data and plasmids can be shared upon reasonable request.

## METHODS

### Constructs, bacteria, and growth conditions

The strains and constructs used in this study are listed in **Table S2**. All bacteria strains used for cloning and expression were grown in LB medium. For the colanic acid secretion assay, the strains were grown in minimal M9 medium supplemented with glucose as a carbon source. *E. coli* JM109 (DE3) Δ*wzabc* strain was generated from *E. coli* JM109 (DE3) by replacing the *wzabc* operon with a chloramphenicol resistance cassette^38^. All expression constructs and JM109 (DE3) strains were verified by DNA sequencing.

### Fluorescence microscopy

*E. coli* BL21 (DE3) strain carrying pET21-Wza-mScarlet3-Wzb-Wzc-sfGFP was grown at 37 °C in LB medium supplemented with 100 mg/ml Ampicillin. The overnight preculture was used to inoculate LB medium supplemented with 100 mg/ml Ampicillin at 30 °C. When the OD_600_ reached 0.6, protein expression was induced with 0.1 mM IPTG for 5 h at 19 °C. The cells were then collected and washed twice with 1 x PBS. The OD_600_ was measured and adjusted to be 0.3 with 1 x PBS. Super-resolution imaging was performed using a LSM980 Airyscan2 confocal microscope (Zeiss). Images were processed using the Zeiss Zen software.

### Protein expression and purification

The *wzabc* operon was amplified from *E. coli* K12 genomic DNA using standard PCR techniques using the primers Wzabc_Fw and Wzabc_Rv (**Table S2**). The vector backbone, excluding the ribosome binding site (RBS), was amplified from the pET21 vector using pET21-V_Fw, and pET21_V_Rv. The construct pET21-Wza-Wzb-Wzc was assembled via Gibson assembly. Subsequently, a Strep-tag was introduced at the C-terminus of Wza using site-directed PCR mutagenesis. The point mutations and pET21-Wzb-Wzc-strep were individually generated using the same mutagenesis method. The fluorescence fusion construct pET21-Wza-mScarlet3-Wzb-Wzc-sfGFP was generated using Gibson assembly.

For large-scale protein purifications, E. coli BL21 Star (DE3) star was transformed with the pET21-Wza-strep-Wzb-Wzc_K540M_ plasmid and grown overnight at 37 °C. An overnight preculture of this transformed strain was used to inoculate 4 liters of LB broth supplemented with 100 µg/ml Ampicillin and grown until an OD600 = 0.8 was reached. Protein expression was induced with 0.1 mM IPTG at 18 °C for 16 h. Cells were harvested by centrifugation at 5000 × g for 15 min and stored at −20 °C. Frozen cells were thawed and suspended in 30 ml lysis buffer (50 mM Tris-HCl pH 8.0, 300 mM NaCl, 20% sucrose, 2 mM MgCl2) supplemented with one tablet cOmplete™ EDTA-free Protease-Inhibitor-Cocktail (Roche). Cells were lysed by six passes through an Emulsiflex-C3 cell disrupter (Avestin). 20% n-dodecyl-ß-D-maltoside (DDM) was directly added to the lysate to a final concentration of 1% and stirred at 4 °C for 2 h. The lysate was ultra-centrifuged at 150,000 x g for 40 min to remove insoluble materials. The supernatant was then incubated at 4 °C overnight with 2 ml Strep-tactin XT resin. The resin was subsequently washed inside a gravity column with 100 ml washing buffer (50 mM Tris-HCl pH 8.0, 150 mM NaCl, 20% sucrose, 6 mM MgCl_2_, 0.05% DDM). Bound protein was eluted as 1 ml fractions with 10 ml total elution buffer (50 mM Tris-HCl pH 8.0, 150 mM NaCl, 20% sucrose, 6 mM MgCl_2_, 50 mM biotin, 0.05% DDM). The protein concentration in the elution fractions was measured by absorbance at 280 nm. Protein-containing fractions were pooled and concentrated using Amicon Ultra 0.5-ml 100-kDa spin concentrators. The concentrated sample was further purified by size-exclusion chromatography using a Superose 6 Increase 10/300 GL column equilibrated with SEC buffer (50 mM Tris-HCl 150 mM NaCl 0.02 % DDM, 5% glycerol, 2 mM MgCl2, 2 mM TCEP). Peak fractions corresponding to the Wza or Wza-Wzc_K540M_ complex were collected separately. The Wza complex was concentrated into 2 mg/ml, and the Wza-Wzc_K540M_ complex was concentrated to 6 mg/ml. Samples were flash frozen in liquid nitrogen (LN) and stored at −80 °C for cryo-EM.

The isolated Wzc_K540M_ octamer was purified from E. coli BL21 Star (DE3) transformed with pET21-Wzb-Wzc_K540M_-strep, following the same procedure described above. After SEC, the peak fraction of Wzc_K540M_ was concentrated to 2.5 mg/ml, flash-frozen in LN, and stored at −80 °C for cryo-EM.

### Structural analysis by cryo-EM

Samples were vitrified on Quantifoil 200-mesh 1.2/1.3 Gold/Copper grids. Briefly, 3.5 μl of the sample was applied onto glow-discharged grids (PELCO easiGlow™ glow discharger, Ted Pella, Inc; glow discharge conditions, -15 mA for 60 s). The grids were plunge-frozen in liquid ethane using a Vitrobot Mark IV (blot time, 4s; blot force, −9; temperature, 6 °C; humidity, 100%).

The vitrified specimens were imaged on a Glacios TEM operating at 200 kV, equipped with a Selectris energy filter set to 10 eV, and a Falcon 4i camera running in counting mode and using Thermo Fisher Scientific EPU software. Automated data acquisition was performed using EPU, and movies were recorded with a nominal magnification of × 130,000, corresponding to a pixel size of 0.91 Å at the specimen level. Movies were recorded with a cumulative electron dose of 40 e-/A2 with a defocus ranging from −2.0 μm to −0.6 μm.

All data-sets were processed in cryoSPARC v4, and cryo-EM density maps were interpreted using ChimeraX 1.9. AlphaFold models were used as initial models, which were iteratively improved with ISOLDE^39^ and PHENIX^40^. Further details are provided in **Table S3.1 and S3.2**.

### Glycan array screening

Glycan array screening was conducted using the Glycan Array 100 kit purchased from RayBioTech (Norcross, GA, USA). This Glycan-100 array was used to assess carbohydrate binding abilities to 100 described glycan structures of Wzc_JR_ and Wzc_Peri_.

Firstly, the biotinylated Wzc_JR_ and Wzc_Peri_ samples were prepared as follows: *E. coli* BL21 Star (DE3) star cells were co-transformed with pACYC-BirA and pET28-Avitag-Wzc_JR_ or pET28-Avitag-Wzc_Peri_ for Wzc_JR_ and Wzc_Peri_ biotinylation *in vivo.* Cells were grown overnight at 37 °C, and then 1 liter LB broth containing 35 µg/ml Kanamycin and 15 µg/ml chloramphenicol was inoculated with preculture then grown to an OD_600_ of 0.8 prior to adding 0.2 mM IPTG and 100 µM biotin, and further grown at 18 °C for 16 h. Cells were collected by centrifugation and resuspended in 50 ml of lysis buffer (50 mM Tris-HCl pH 7.5, 500 mM NaCl, 10 mM imidazole, 10% glycerol). After adding 1 tablet of protease inhibitor cocktail (11873580001, Roche), cells were lysed by sonication (BANDELIN SONOPLUS; Amplitude 58%; 1 sec on; 8 sec off, 35 min total). After soluble fractions were obtained, batch purification was conducted with 3 ml (bed volume) nickel resin. After 1 h incubation at 4 °C, Nickel resin was washed with 200 ml wash buffer (50 mM Tris-HCl pH 7.5, 500 NaCl, 20 mM imidazole, 10% glycerol), and bound proteins were eluted in 15 ml (50 mM Tris-HCl pH 7.5, 500 mM NaCl, 300 mM imidazole, 10% glycerol). Elution fractions were concentrated using 10 kD cut-off Amicon® Ultra Centrifugal Filters (Merk), and then loaded onto Superdex 200 increase 100/30 GL gel filtration column equilibrated with SEC running buffer (50 mM Tris-HCl pH 7.5, 150 NaCl, 5% glycerol). The peak elution fractions were first analysed on SDS-PAGE and then pooled, concentrated, aliquoted, flash frozen, and stored in -80 °C.

The glycan screening assay was performed according to the manufacturer’s protocols. The biotinylated recombinant proteins at three different concentrations (Wzc_JR_, 25 µg/ml, 50 µg/ml, and 100 µg/ml; Wzc_Peri_, 50 µg/ml, 100 µg/ml, and 200 µg/ml), were added to array wells, with one buffer-only control well. The samples were incubated overnight with gentle rocking. Glycan-protein binding was detected by incubation with Cy3 equivalent dye-conjugated streptavidin for 1 h at 4 °C. The washing procedure was strictly performed according to the manufacturer’s protocols. The glass slide was dried with nitrogen flow before storage.

The fluorescent signals were scanned using a GenePix 4300 microarray scanner (Molecular Devices) with excitation at 532 nm and emission at 568 nm. Signal intensities were quantified using GenePix Pro 7 software (7.3.0.0). Background correction was performed by subtracting signal values obtained from buffer-only controls to reduce noise. Data were subsequently normalized following the manufacturer’s guidelines (GA-Glycan-100-SW). To ensure data quality, four outlier signals (out of 2400 data points) inconsistent with technical replicates were excluded as likely artifacts. Final data visualization, including heatmap generation, was performed using RStudio (version 2024.04.2+764).

### Colanic acid secretion assay

The colanic acid secretion assay was performed according to previous studies with small modifications^23,24,41^. Briefly, colanic acid was precipitated from spent M9 culture after removing bacterial cells by adding three equivalent volumes of acetone. Added Cys·HCl reacted with fucose to generate a coloured product. The concentration of this product was measured colourimetrically to determine the relative concentration of fucose as a proxy for colanic acid concentration.

E. *coli* JM109 (DE3) Δ*wzabc* strain was made from *E. coli* JM109 (DE3) using λred recombination and chloramphenicol resistance cassette as a selection marker^38^. *E. coli* JM109 (DE3) Δ*wzabc* cells were transformed with pET21-Wza-Wzb-Wzc-strep and the derivatives. Pre-cultures were grown in LB supplemented with appropriate antibiotics at 37 °C overnight. Then the pre-cultures were diluted (1:50) into 1 x M9 media supplemented with 0.4% glucose as a carbon source, and appropriate antibiotics. The cultures were further grown at 19 °C for 1 day to reach around OD_600_= 0.6. Then 0.1 mM IPTG were added to each culture, and cultures were further grown for 24 h. The OD_600_ was measured for all cultures, and the volume of the cultures were normalised based on OD_600_. The cell pellets were discarded after centrifugation at 20,000 x g for 30 min. 4 x 400µl from each sample were taken and transferred into 4 EP tubes and 100 µl TCA was added to each tube to precipitate the protein for 4 h. The protein pellets were discarded after centrifugation at 20,000 x g for 30 min, and 450 µl was transferred into a new 2-ml EP tube and 1350µl acetone was added to precipitate colanic acid overnight. The secreted colanic acid was spun down by centrifugation at 20,000 x g for 60 min, and dried for 2 h using a vacuum concentrator (RVC 2-18 CDplus, Avantor). The purified samples were resuspended in 50 μl Milli-Q H_2_O. After adding 200 μl of H_2_SO_4_/H_2_O (6:1 v/v), the samples were then heated up to 95 °C for 30 min and then cooled to room temperature. The absorbances of each sample (200 μl) were measured at both 396 and 427 nm within a 96-well U shape plate using a microplate reader (TECAN SPARK Microplate Reader, TECAN). The difference between OD_390_ and OD_427_ is marked as A1. Afterwards, 6 μl freshly prepared cysteine hydrochloride (Cys·HCl, 3% (w/v)) solution was added into each sample well and mixed by pipetting. The absorbances of each sample were measured again at both 396 and 427 nm. The current difference between OD_390_ and OD_427_ is marked as A2. The difference between A2 and A1 was then plotted as a violin plot using graphpad prism 10.

### Mass spectrometry

For proteomic analysis, purified wild type Wzc from SEC was digested using trypsin. For that, 1 μg of protein was solubilized in 100 μl 500 mM TEAB (pH 8) followed by one hour reduction with final concentration of 5 mM TCEP at 56 °C, and 30 minutes alkylation with a final concentration of 10 mM MMTS at room temperature. Trypsin (Promega) was then added at a ratio of 0.05 μg Trypsin per 1 μg protein, and digestion was carried out overnight at 37 °C while shaking at 800 rpm.

Peptides were directly subjected to desalting using the Bravo Automated Liquid Handling Platform (Agilent) applying the standard peptide clean-up method version 3.0 (protein sample prep workbench v.3.2.0) and using reversed-phase S cartridges (G5496-60033, Agilent).

Next, desalted peptides were directly used for phosphopeptide enrichment. Enrichment was conducted on the Bravo system using the standard phosphopeptide enrichment method version 2.1 using Fe(III)-NTA cartridges (G5496-60085, Agilent).

For LC-MS analyses peptides were vacuum dried, resuspended in 3 μl of 0.1% formic acid (FA) in water, sonicated for 5 minutes, and transferred to HPLC vials. LC-MS/MS analyses were performed using a Dionex Ultimate 3000 n-RSLC system (Thermo Fisher Scientific) coupled to an Orbitrap Exploris Hybrid mass spectrometer (Thermo Fisher Scientific). One nanogram of peptides was loaded onto a 50 cm Low-Load μPAC Neo HPLC analytical column (Thermo Fisher Scientific). The peptides were separated at a flow rate of 500 nl/min with a nonlinear gradient over 5. 5 minutes from 99% MS buffer A (0.1% FA in water) to 10% MS buffer B (0.1% FA in 80% ACN), followed by 10 minutes to 35% buffer B and finally 4 minutes to 98% buffer B at 125 nl/min.

Ionization was achieved via electrospray using a glass emitter (Bruker). Data acquisition was performed using SII software within the Xcalibur suite (v4.4.16.14). The instrument operated in “top speed” mode with a 1.5 s cycle for MS/MS acquisition of doubly and triply charged peptides. Fragmentation was conducted using higher-energy collisional dissociation (HCD), and peptides were measured in the Orbitrap (HCD/OT).

Raw data was analysed using MaxQuant (v2.6.7.0) against the *Escherichia coli* database (strain K12, downloaded 2025-03-28 from Uniprot). Search parameters included: trypsin as enzyme allowing two missed cleavages; maximum modifications on one peptide: 8; fixed modification: β-methylthiolation on cysteine (+45.98 Da); variable modifications: oxidation on methionine (+15.99 Da) and phosphorylation at serine, threonine, and tyrosine (+79.97 Da); search peptide tolerance: 4.5 ppm; false discovery rate (FDR): 1%. All peptides were used for quantification. Removal of contaminants, missing values and phosphorylated sites with localization probability < 85% was done with support of Perseus (v2.1.1.0). Summed intensities of phosphorylation site combinations were calculated based on Peptide Spectrum Match (PSM) representing confirmed identification of peptide sequences. Association analysis (frequent itemset mining approach) was performed in R to evaluate the co-occurrence of phopshorylations (itemsets). Support is defined as the summed intensity of observed modifications normalized to total measured intensities. Lift quantifies deviation from statistical independence, with values >1 indicating co-enrichment, <1 suggesting anti-co-occurrence, and 1 reflecting expected random association. Mass spectrometry data have been deposited to the ProteomeXchange Consortium via the PRIDE partner repository (dataset and identifier on request)^42^.

### Protein structure prediction

Protein structures were predicted with a local installation of AlphaFold 3^29^. The protein sequences of putative colanic acid polymerase Wzy (gene name *wcaD*; uniprot ID, P71238), and the co-polymerase Wzc (gene name *wzc*; uniprot ID, P76387) were used for structural predictions.

