## Supplemental file for "Molecular insights into the capsular polysaccharide transporter Wza-Wzc complex"

### SUPPLEMENTARY FIGURES

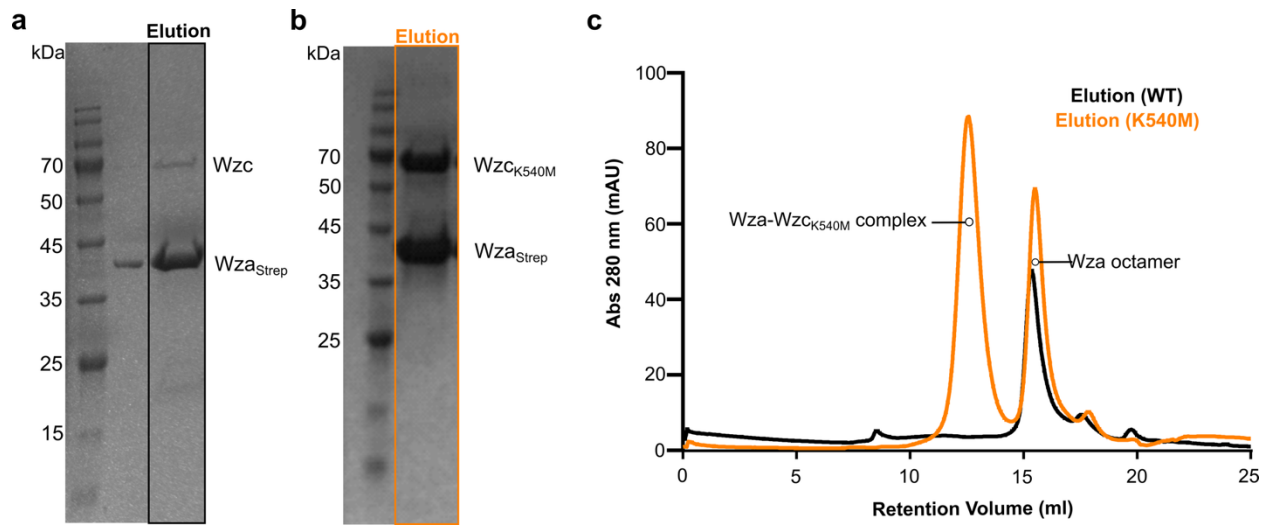

**Fig. S1. Purification of the Wza-Wzc complex.** (a) A little amount of wild-type Wzc was co-purified with Wza<sub>Strep</sub>. SDS-PAGE analysis shows the elution profile of *E. coli* BL21 Star (DE3) cells overexpressing the full *wzabc* operon with C-terminal Strep-tag on Wza from Strep-tactin XT resin. (b) The octameric variant Wzc<sub>K540M</sub> was co-purified with Wza<sub>Strep</sub>. SDS-PAGE analysis of elution from *E. coli* BL21 Star (DE3) expressing the full *wzabc* operon with C-terminally Strep-tagged Wza and K540M variant of Wzc. (c) Size-exclusion chromatography (SEC) profiles of the elution samples from panels a and b. The Wza-Wzc<sub>K540M</sub> complex purified by SEC was used for subsequent cryo-EM analysis.

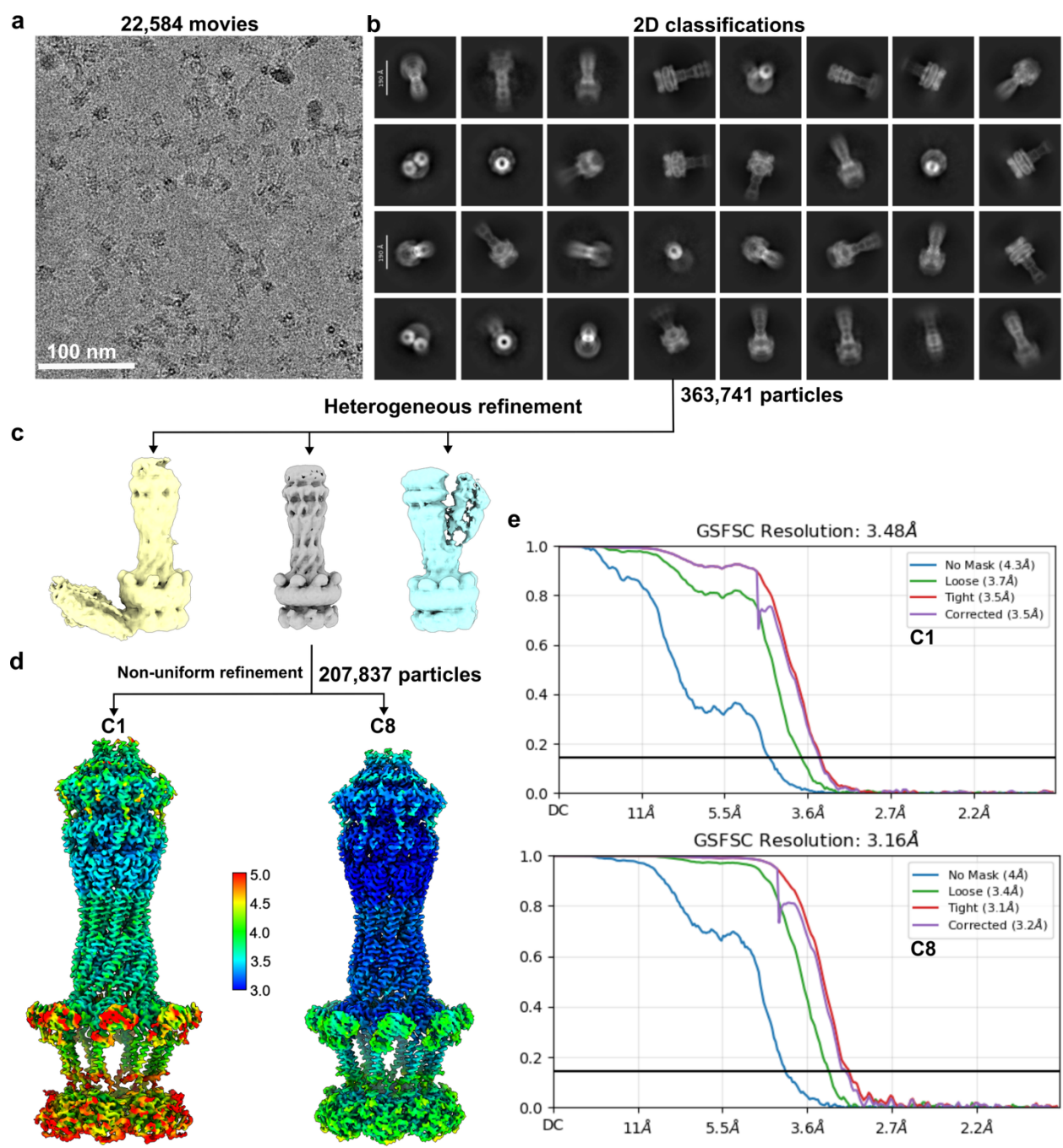

**Fig. S2. Cryo-EM data processing workflow for the Wza-Wzc<sub>K540M</sub> complex.** (a) Representative cryo-EM micrograph of the Wza-Wzc<sub>K540M</sub> complex. Scale bar: 100 nm. (b) Selected 2D class averages used for subsequent analysis. Scale bar: 190 Å. (c) 3D classification step to exclude poor-quality or junk particles. (d) Final cryo-EM density maps generated using non-uniform refinement under C1 and C8 symmetries, colored according to local resolution. (e)

20 Gold standard Fourier Shell Correlation (FSC) curves for the refined C1 (top panel) and C8
21 (bottom) cryo-EM maps.
22
23

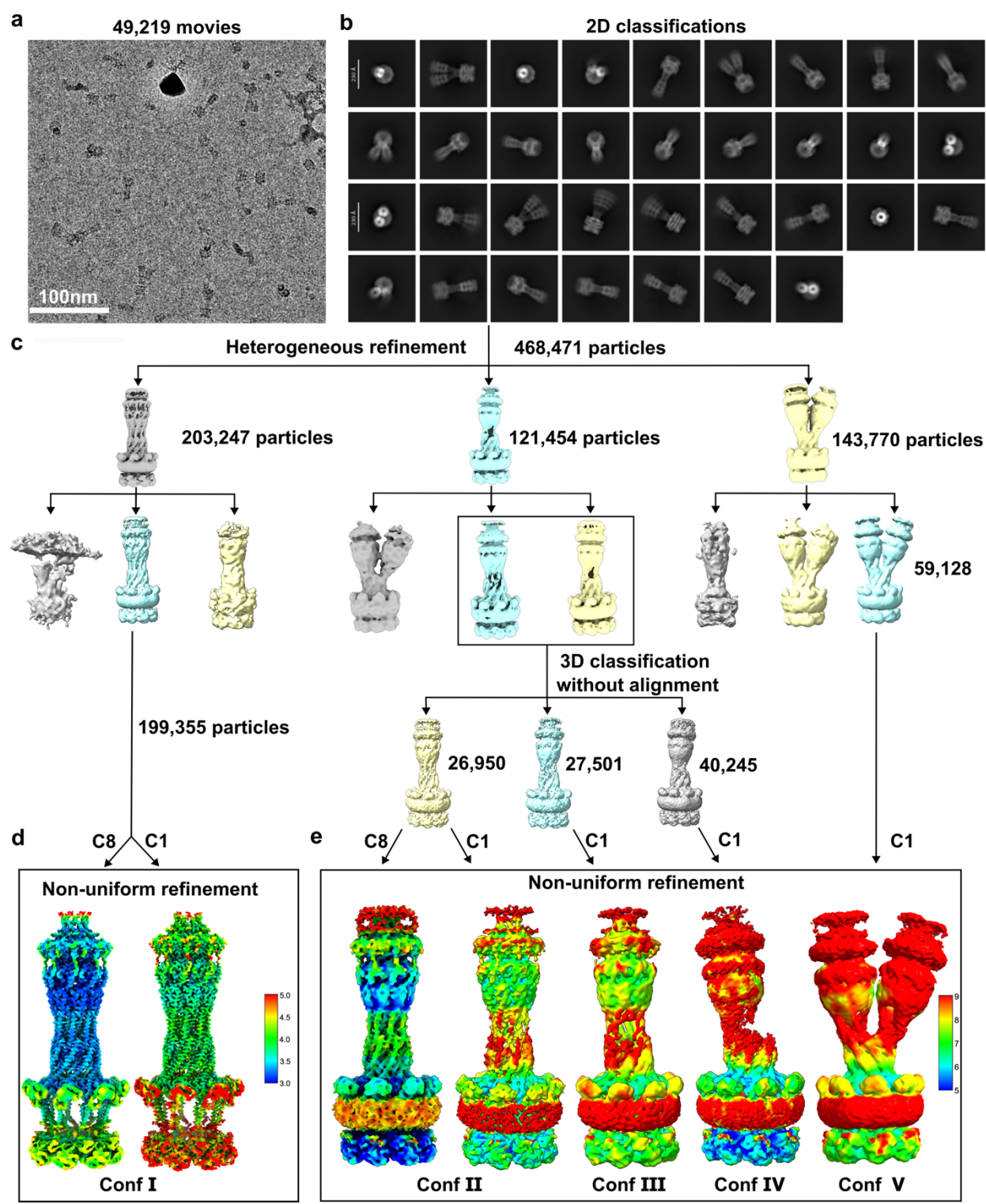

**Fig. S3. Data processing of the EDTA-treated Wza-Wzc<sub>K540M</sub> complex.** (a) Representative cryo-EM micrograph of the Wza-Wzc<sub>K540M</sub> complex following treatment with 10 mM EDTA. Scale bar: 100 nm. (b) Selected 2D class averages used for subsequent 3D classification. Scale bar:

28 230 Å **(c)** Workflow involving two rounds of 3D classification, non-uniform refinement, and
29 alignment-free 3D classification to resolve multiple conformational states of the complex. **(d-e)**
30 Five distinct conformational states (Confs I – V) were identified, with corresponding density maps
31 colored according to local resolution.
32
33
34

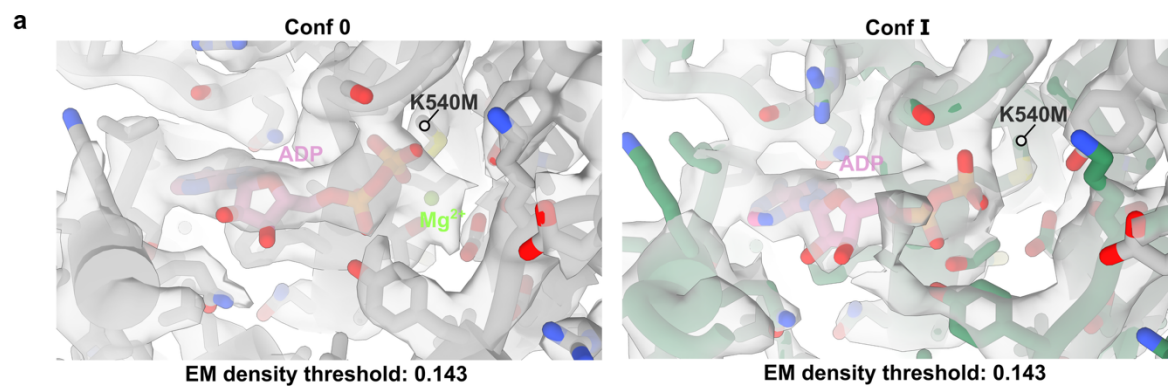

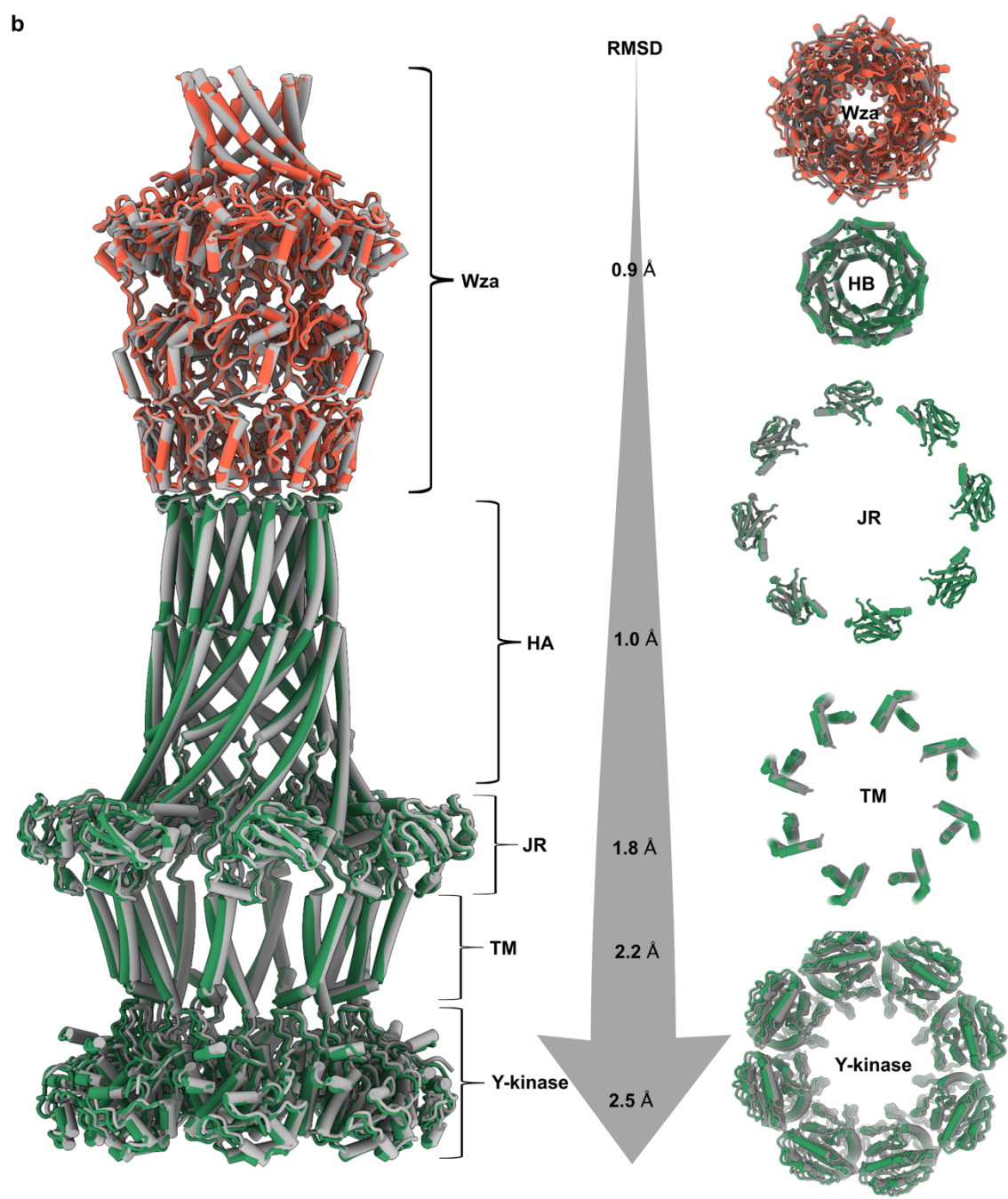

**Fig. S4 Structural comparison between Conf I and Conf 0 of the Wza-Wzc complex. (a)**

Cryo-EM maps of the ADP-Mg<sup>2+</sup> binding pockets in the Wza-Wzc<sub>K540M</sub> complex. The Conf 0 structure (shown in **Fig. 1c**) is colored grey while the Conf I structure (shown in **Fig. 3b**) is overlaid for comparison. **(b)** Structural superposition of Conf I and Conf 0 of the Wza-Wzc<sub>K540M</sub> complex aligned on the Wza translocon, including local RMSD values and cross-sectional views highlighting differences in Wza and the individual subdomains of Wzc.

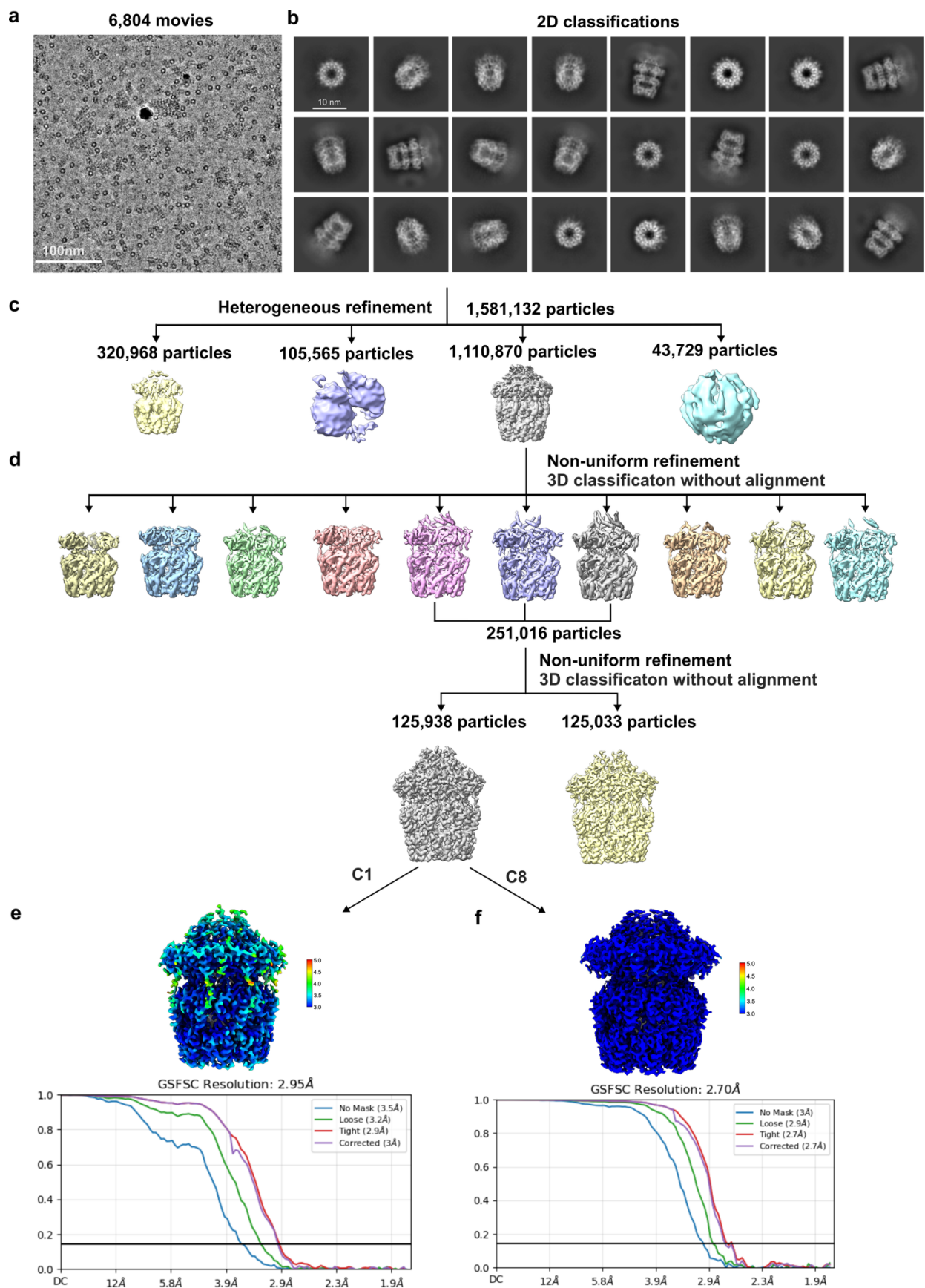

**Fig. S5. Data processing workflow for the Wza complex.** (a) Representative cryo-EM micrograph of the Wza complex. (b) Selected high-quality 2D class averages used for further analysis. (c) Initial 3D classification to eliminate junk and broken particles. (d) Two rounds of non-uniform refinement and alignment-free 3D classification to further remove broken particles. (e) Final cryo-EM map and corresponding FSC curve obtained from non-uniform refinement under C1 symmetry. (f) Cryo-EM map and FSC curve obtained from non-uniform refinement with C8 symmetry.

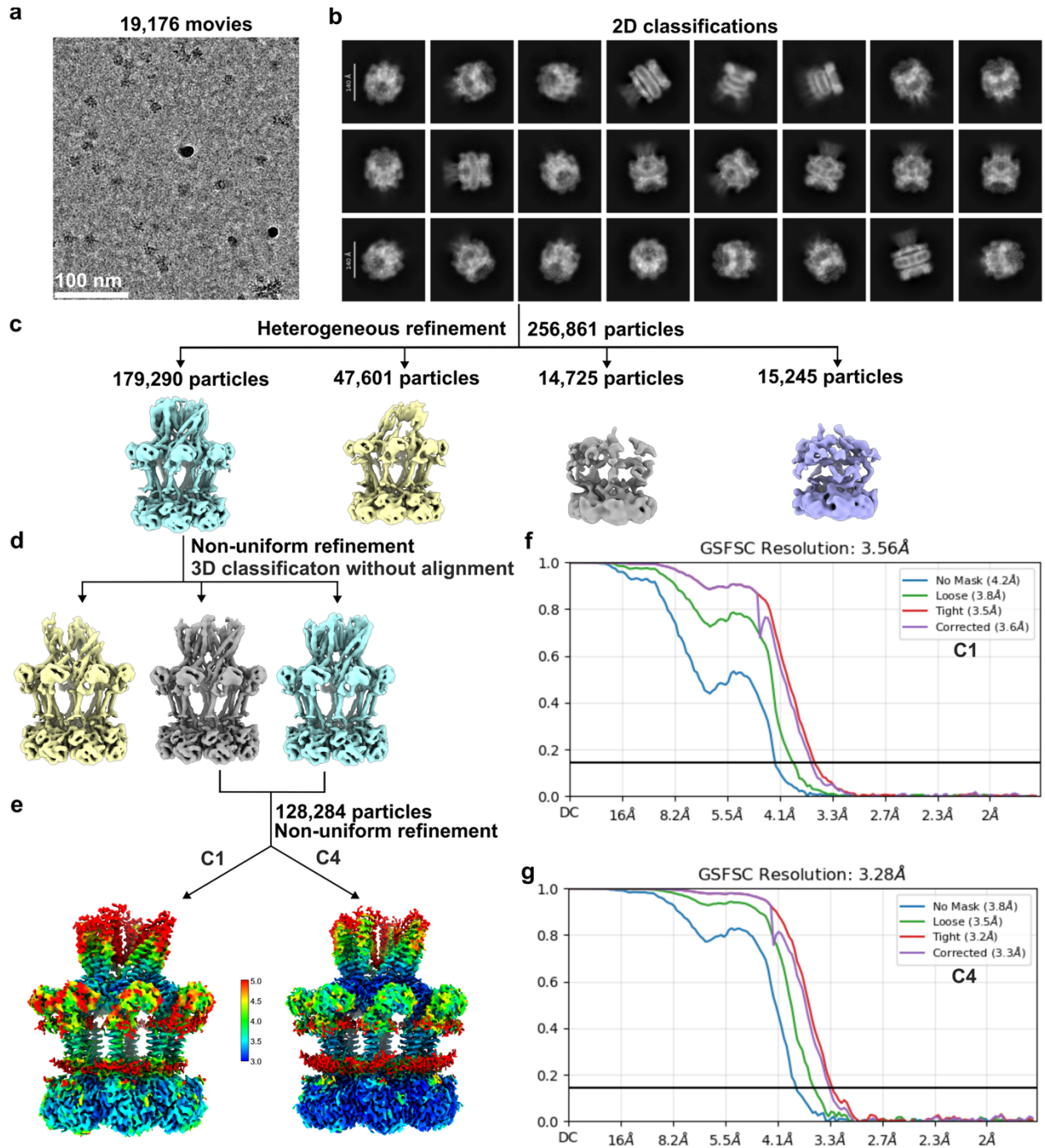

**Fig. S6. Data processing workflow for the Wzc<sub>K540M</sub> complex.** (a) Representative cryo-EM micrograph of the Wzc<sub>K540M</sub> complex. (b) Selected high-quality 2D class averages from 2D classifications. (c) 3D classification step used to remove junk classes. (d) One round of non-uniform refinement followed by alignment-free 3D classification to eliminate remaining low-quality classes. (e) Final cryo-EM density maps obtained from non-uniform refinements with C1 and C4

symmetries applied. (f) FSC curve for the C1-refined map. (g) FSC curve corresponding to the C4-refined map.

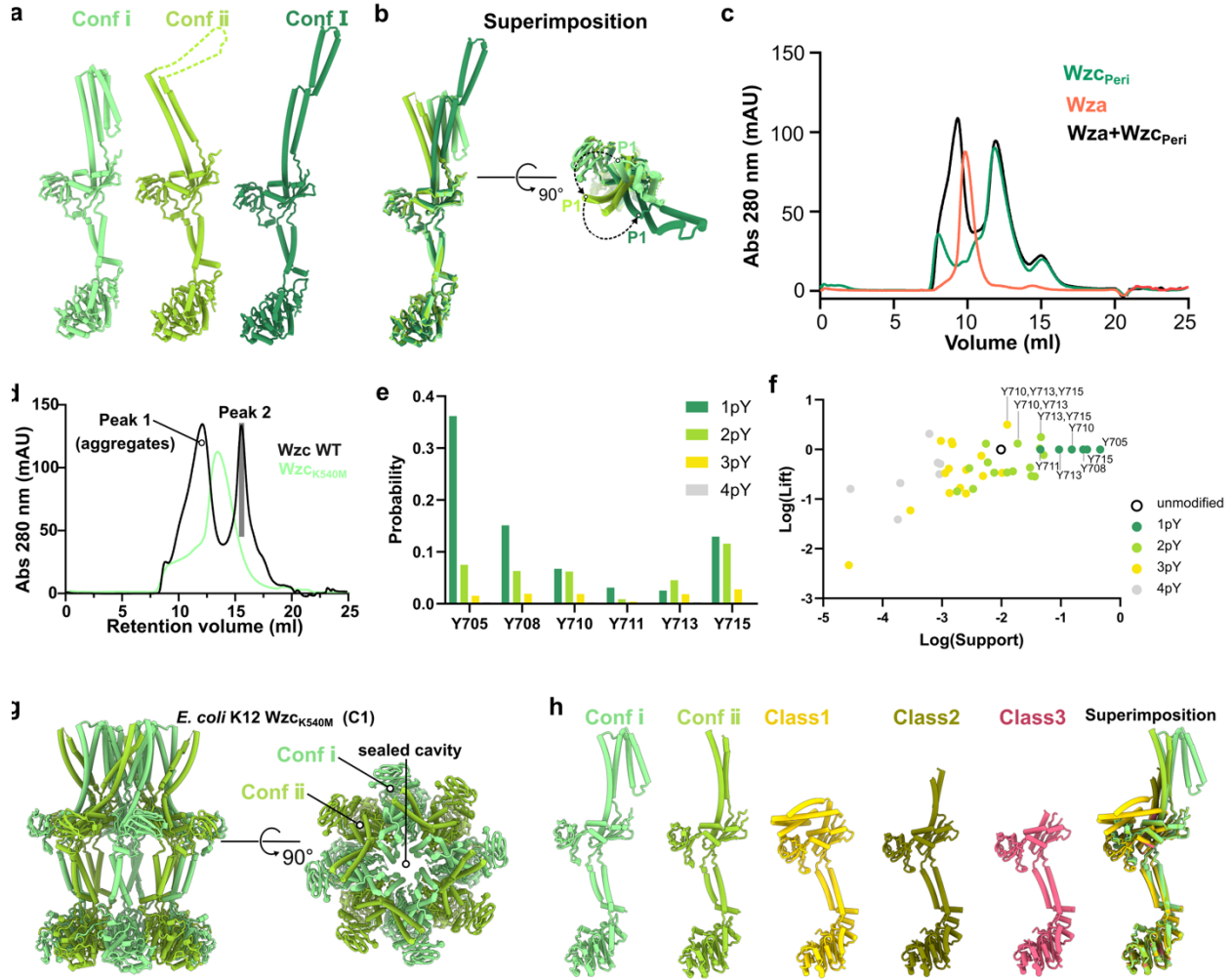

**Fig. S7. Structural comparison between *E. coli* K12 and K30 $Wzc_{K540M}$ structures. (a)** Structural comparison of protomers of the Wzc octamer and the Wza-Wzc<sub>K540M</sub> complex. (b) Superimposition of these protomers indicating the trajectory of HA to recruit the Wza translocon. (c) SEC profiles of Wzc<sub>Peri</sub>, Wza alone, and a combined sample of Wzc<sub>Peri</sub> and Wza, with the mixture containing equal amounts of each protein as used separately. (d) Size-exclusion chromatography (SEC) profiles of purified wild type Wzc and Wzc<sub>K540M</sub>. Peak 1 sample of wild-

type Wzc corresponds to soluble aggregates, while Peak 2 represents a lower-order oligomeric species. Separation was performed using a Superose 6 Increase 10/300 GL column following affinity purification with a Strep-Tactin-XT column. **(e)** Phosphorylation probability of each tyrosine position at Y-tails with 1pY, 2pY, 3pY, and 4pY. **(f)** Association analysis of different phosphorylated sites at Y-tails phosphorylation sites indicating the prevalence (Support) and occurrence (Lift) of single or higher order phosphorylations. **(g)** Cryo-EM structure of *E. coli* K12 Wzc<sub>K540M</sub> (C1). **(h)** Structural comparison of different states of the protomers of *E. coli* K12 and K30 Wzc<sub>K540M</sub> structures<sup>1</sup>. Conf i , Wzc protomer at Conf i state from the Wzc<sub>K540M</sub> octamer; Conf ii , Wzc protomer at Conf ii state from Wzc<sub>K540M</sub> octamer; Conf I, Wzc protomer from the Conf I Wza-Wzc<sub>K540M</sub> complex. Superposition of these Wzc protomers showing HA under large conformational changes.

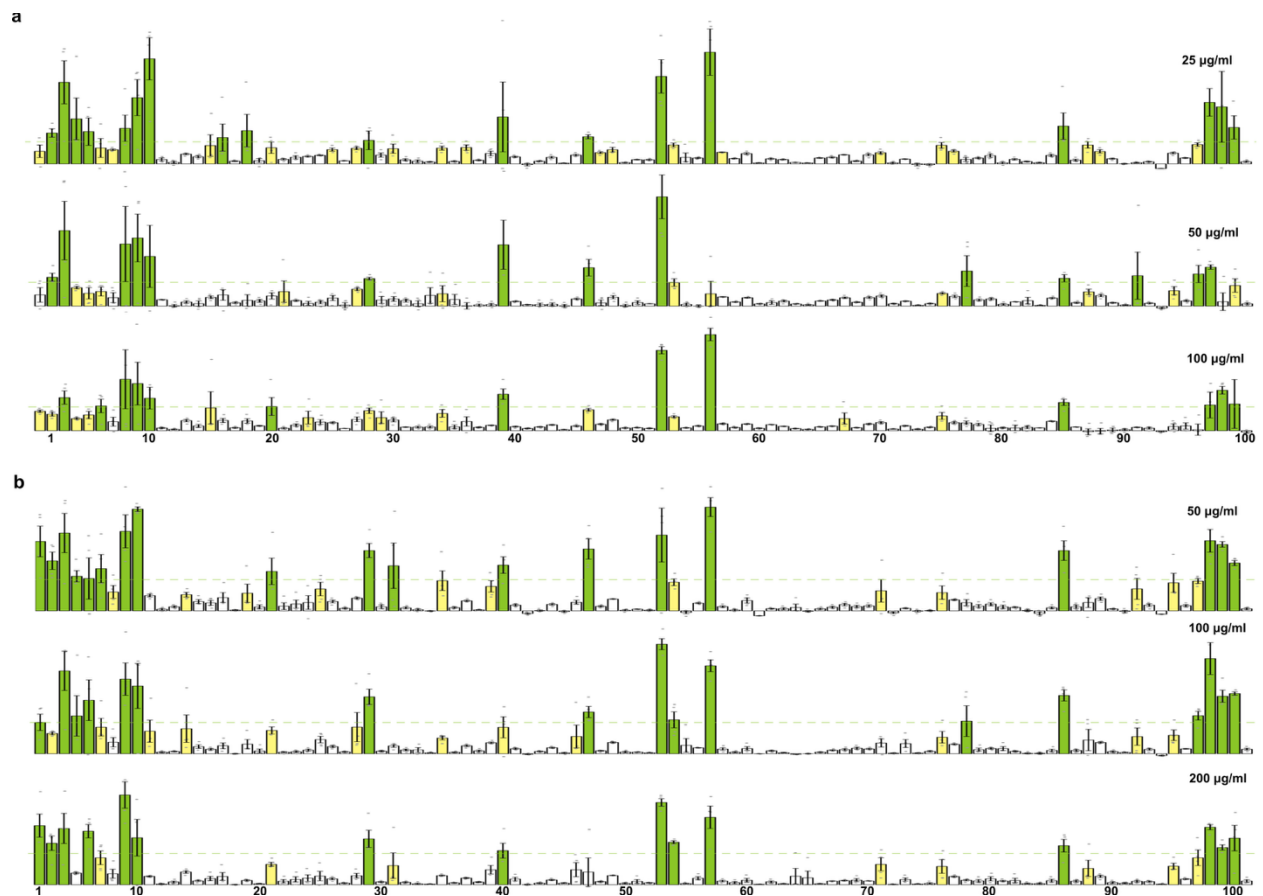

**Fig. S8. Glycan array 100 screening for the JR and periplasmic domains of Wzc. (a-b)**

Glycan binding profiles of Wzc<sub>JR</sub> (a) and Wzc<sub>Peri</sub> (b) at three different concentrations, as labels for each panel. Glycan-binding profile of biotinylated proteins detected using Cy3-conjugated streptavidin on the Glycan 100 array. The Y-axis represents relative fluorescence intensity, indicating the extent of protein binding to each glycan. The X-axis corresponds to glycan ID numbers as defined in the Glycan 100 array. The bars are colored according to the values of the relative fluorescence intensity (intensity > 1000, green; 500 < intensity < 1000, yellow; intensity < 500, white).

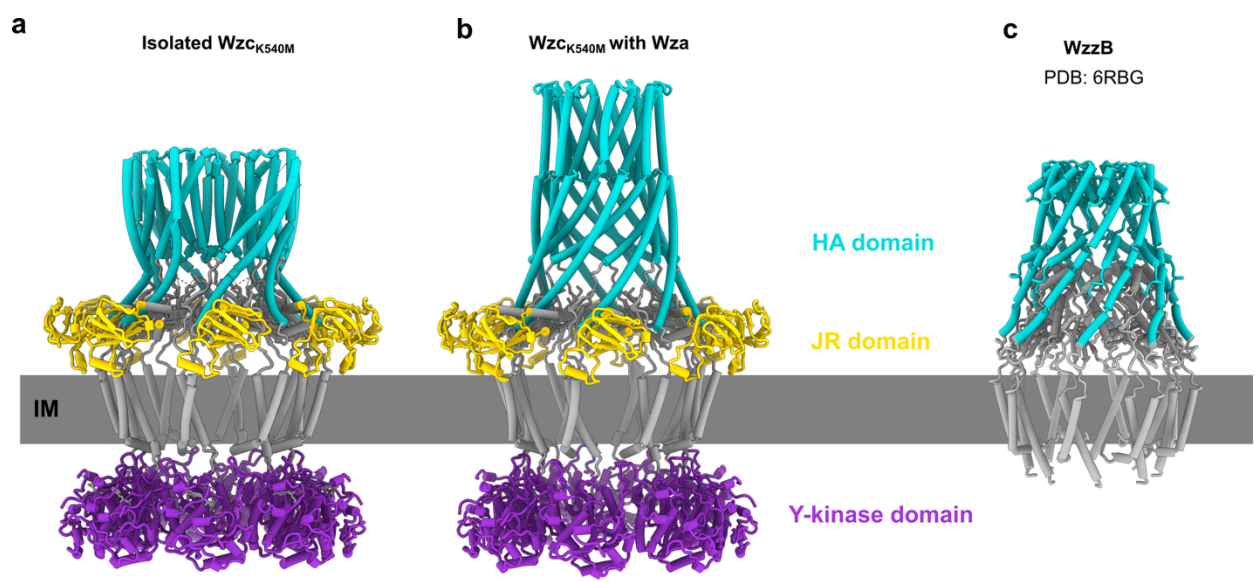

**Fig. S9 Structural comparison between Wzc and WzzB. (a)** Model of the isolated Wzc<sub>K540M</sub> octamer. **(b)** Model of the Wzc<sub>K540M</sub> when bound to the Wza translocon. **(c)** Model of the full-length WzzB octamer<sup>2</sup>.

111 **SUPPLEMENTARY TABLES**

112 **Table S1. Mass spectrometry analysis of the phosphorylation pattern on the Y- tail of Wzc.**

| Peptide sequence with Phospho<br>(STY) Probabilities | PEP | Score | Intensity | Phosphorylation sites |
| --- | --- | --- | --- | --- |
| ASAY(1)QDY(1)GY(1)YEYKY | 0 | 319,37 | 5965600 | Y705, Y708, 710Y |
| ASAY(1)QDY(0.027)GY(0.472)Y(0.472)<br>EY(0.009)EY(0.019)KSDAK | 0 | 251,75 | 1,32E+08 | Y705 |
| ASAYQDYG(0.034)Y(0.966)EY(1)EY(0<br>.988)KS(0.013)DAK | 0 | 301,03 | 2023000 | YY08, Y710, Y711 |
| ASAYQDYG(0.026)Y(0.974)EY(0.995)<br>EY(0.004)KSDAK | 0 | 325,11 | 5682000 | Y708, Y710 |
| ASAY(0.006)QDY(0.994)GY(0.067)Y(0.<br>933)EYKY | 0 | 275,17 | 9026500 | Y708, Y711 |
| ASAYQDY(0.031)GY(0.857)Y(0.109)EY(<br>0.079)EY(0.902)KS(0.022)DAK | 0 | 254,83 | 84666000 | Y708, Y713 |
| ASAYQDY(0.005)GY(0.994)Y(0.001)EY<br>EYK | 0 | 283 | 52737000 | Y708 |
| RAS(0.648)AY(0.352)QDYG(0.137)Y(0<br>.863)EY(0.027)EY(0.965)KS(0.007)DAK | 0 | 266,78 | 1142400 | Y710, Y713 |
| RAS(0.622)AY(0.353)QDY(0.025)GY(0.<br>858)Y(0.141)EY(0.002)EY(0.996)KS(0.0<br>03)DAK | 0 | 240,82 | 4314600 | Y710, Y715 |
| ASAY(0.001)QDY(0.826)GY(0.155)Y(0.<br>062)EY(0.962)EY(0.967)KS(0.026)DAK | 0 | 259,82 | 29923000 | Y713, Y715 |
| RAS(0.427)AY(0.57)QDY(0.14)GY(0.83<br>9)Y(0.028)EY(0.028)EY(0.967)KS(0.001<br>)DAK | 0 | 254,16 | 3360500 | Y715 |
| ASAY(1)QDY(1)GYEY(0.033)EY(0.967<br>)K | 0 | 258,53 | 3693200 | Y705, Y708, Y711 |
| AS(0.038)AY(0.962)QDY(1)GY(0.295)Y(<br>0.705)EY(0.049)EY(0.946)KS(0.005)DA<br>K | 0 | 270,75 | 1817000 | Y705, Y708, Y715 |
| ASAY(1)QDYG(1)YEEYK | 0 | 310,01 | 23300000 | Y705, Y708 |
| ASAY(1)QDYGYYEY(0.028)EY(0.972)K | 0 | 364,34 | 19400000 | Y705, Y710 |
| RASAYQDYG(0.937)Y(0.063)EY(0.00<br>5)EY(0.995)KS(0.001)DAK | 0 | 252,44 | 5106400 | Y705, Y711 |
| ASAYQDYG(0.757)Y(0.298)EY(0.917)<br>EY(0.027)KS(0.001)DAK | 3,4E-283 | 235,85 | 5315800 | Y710 |
| RAS(0.219)AY(0.781)QDY(1)GY(0.984)<br>Y(0.016)EY(0.99)KS(0.01)DAK | 7,5E-280 | 245,66 | 1511900 | Y708, Y710, Y713 |
| RAS(0.22)AY(0.782)QDY(0.999)GY(0.9<br>84)Y(0.016)EY(0.001)EY(0.989)KS(0.01<br>)DAK | 7,5E-280 | 245,66 | 1784500 | Y708, Y710, Y715 |
| RASAY(0.001)QDY(0.982)GY(0.015)Y(0<br>.001)EY(0.001)EY(0.988)KS(0.01)DAK | 7,3E-255 | 238,79 | 4371000 | Y708, Y715 |
| ASAY(1)QDY(0.999)GY(0.01)Y(0.135)E<br>Y(0.857)EY(0.956)KS(0.044)DAK | 2,6E-254 | 230,27 | 5329800 | Y705, Y708, Y713, Y715 |
| ASAY(0.01)QDY(0.989)GY(0.924)Y(0.0<br>8)EY(0.996)EY(0.973)KS(0.027)DAK | 5,2E-228 | 229,86 | 1246900 | Y708, Y710, Y713, Y715 |
| ASAYQDY(0.006)GY(0.609)Y(0.391)EY(<br>0.993)EY(0.969)KS(0.032)DAK | 1,7E-227 | 214 | 41992000 | Y711, Y713 |
| ASAY(0.001)QDY(0.98)GY(0.228)Y(0.8<br>04)EY(0.973)EY(0.954)KS(0.059)DAK | 2,8E-227 | 223,44 | 7317600 | Y708, Y713, Y715 |

|  |  |  |  |  |
| --- | --- | --- | --- | --- |
| RAS(0.051)AY(0.933)QDY(0.016)GY(0.003)Y(0.008)EY(0.982)EY(0.008)KSDA | 1,9E-202 | 218,72 | 2389500 | Y705, Y713 |
| K |  |  |  |  |
| ASAYQDY(0.01)GY(0.88)Y(0.153)EY(0.961)EY(0.963)KS(0.033)DAK | 3E-202 | 217,89 | 33255000 | Y708, Y711, Y713 |
| ASAYQDY(0.992)GY(0.031)Y(0.979)EY(0.89)EY(0.105)KS(0.004)DAK | 4E-202 | 215,83 | 9139900 | Y705, Y710, Y711 |
| ASAY(0.973)QDY(0.029)GY(0.988)Y(0.016)EY(0.995)EY(0.963)KS(0.035)DAK | 1,2E-178 | 203,5 | 5857500 | Y705, Y710, Y713, Y715 |
| RAS(0.05)AY(0.879)QDY(0.547)GY(0.53)Y(0.487)EY(0.507)EY(0.938)KS(0.062 | 2,5E-175 | 209,01 | 2687600 | Y705, Y715 |
| )DAK |  |  |  |  |
| ASAYQDY(0.845)GY(0.153)Y(0.002)EY(0.016)EY(0.958)KS(0.026)DAK | 1,5E-157 | 203,23 | 85133000 | Y713 |
| ASAY(1)QDY(1)GY(0.986)Y(0.028)EY(0.979)EY(0.006)KS(0.001)DAK | 2,8E-157 | 197,99 | 3648300 | Y705, Y708, Y710, Y713 |
| RAS(0.069)AY(0.917)QDY(0.016)GY(0.999)Y(0.999)EY(0.967)EY(0.033)KS(0.0 | 1,7E-133 | 189,51 | 1162200 | Y705, Y710, Y711, Y713 |
| 01)DAK |  |  |  |  |
| AS(0.099)AY(0.9)QDY(0.003)GY(0.933)Y(0.064)EY(0.005)EY(0.987)KS(0.008)D | 8,4E-119 | 175,6 | 1193300 | Y705, Y710, Y715 |
| AK |  |  |  |  |
| ASAYQDY(0.018)GY(0.036)Y(0.237)EY(0.892)EY(0.782)KS(0.035)DAK | 3,9E-103 | 163,61 | 51991000 | Y711 |
| AS(0.083)AY(0.915)QDY(0.001)GY(0.002)Y(0.04)EY(0.959)EY(0.986)KS(0.013) | 2,22E-88 | 166,09 | 343190 | Y705, Y713, Y715 |
| DAK |  |  |  |  |

|  |  |  |  |  |
| --- | --- | --- | --- | --- |
| AS(0.006)AY(0.994)QDY(1)GY(0.003) | 8,46 | 169,09 | 3002100 | Y705, Y708, Y713 |
| EY(0.02)EY(0.977)K | E-87 |  |  |  |
| AS(0.002)AY(0.998)QDY(0.046)GY(0.90 | 9,77 | 158,06 | 314100 | Y705, Y710, Y713 |
| 9)Y(0.065)EY(0.969)EY(0.011)K | E-72 |  |  |  |
| AS(0.003)AY(0.433)QDY(0.549)GY(0.17 | 5,64 | 131,94 | 3715600 | Y711, Y713, Y715 |
| 1)Y(0.883)EY(0.958)EY(0.961)KS(0.043 | E-49 |  |  |  |
| )DAK |  |  |  |  |
| RAS(0.1)AY(0.309)QDY(0.602)GY(0.98 | 2,51 | 130,05 | 343500 | Y710, Y711 |
| 9)Y(0.993)EY(0.742)EY(0.253)KS(0.012 | E-36 |  |  |  |
| )DAK |  |  |  |  |
| RAS(0.23)AY(0.751)QDY(0.06)GY(0.96 | 6,16 | 128,67 | 1198500 | Y710, Y711, Y713 |
| 1)Y(0.994)EY(0.861)EY(0.137)KS(0.006 | E-36 |  |  |  |
| )DAK |  |  |  |  |
| AS(0.005)AY(0.995)QDY(0.99)GY(0.982 | 3,47 | 99,86 | 1130600 | Y705, Y708, Y710, Y715 |
| )Y(0.808)EY(0.239)EY(0.887)KS(0.094) | E-17 |  |  |  |
| DAK |  |  |  |  |
| RAS(0.185)AY(0.359)QDY(0.719)GY(0. | 4,82 | 77,045 | 181200 | Y710, Y711, Y713, Y715 |
| 873)Y(0.897)EY(0.946)EY(0.906)KS(0.1 | E-10 |  |  |  |
| 15)DAK |  |  |  |  |
| AS(0.052)AY(0.947)QDY(0.798)GY(0.62 | 1,09 | 49,528 | 169440 | Y705, Y711, Y715 |
| 9)Y(0.851)EY(0.731)EY(0.852)KS(0.139 | E-05 |  |  |  |
| )DAK |  |  |  |  |

113

114

115

116

117

118

| Primer name | Sequence (5' → 3') | Products |
| --- | --- | --- |
| Wzabc_Fw | AACAATTCCCCTCTAGAAATAATTAATAGTGCACAGGATA<br>ATTACTCTGCC | pET21-Wza-Wzb-Wzc |
| Wzabc_Rv | CAGTGGTGGTGGTGGTGGTGTTTCGCATCCGACTTATAT<br>TGTATTCGTA |  |
| pET21-V_Rv | TATTTCTAGAGGGGAATTGTTATCCGCTCA |  |
| pET21_V_Fw | CACCACCACCACCACCACTG |  |
| Wza-strep_Fw | TGGGAGAACTTGTACTTCCAGAGCTGGAGCCACCCGCA<br>GTTCGAAAAATGAACGGATACAGCCAGCGACAT | pET21-Wza-strep-Wzb-Wzc |
| Wza-strep_Rv | AAGTACAAGTTCTCCCAATTATGAATATCAGACGCCGTAT<br>CGGTCATGTAACGGACACCGCTAATAG |  |
| WzcK540M_Fw | AGCCCGTCAATTGGTATGACCTTTGTCTGCGCC | Any<br>constructs<br>containing<br>K540M<br>mutation |
| WzcK540M_Rv | GGCGCAGACAAAGGTCATACCAATTGACGGGCT |  |
| Wzc_ΔJR_Fw | GGTGGCGGTGGCTCGTACTCCACGCTGGGGATGATC | pET21-Wza-Wzb-Wzc<sub>ΔJR</sub>-<br>strep |
| Wzc_ΔJR_Rv | ACGAGCCACCGCCACCAATATCGAGGTCGAGATCGTCC<br>A |  |
| WzcR374E_Fw | ATTGTCgaaCTGACCCGCGATGTCGAG | pET21-Wza-Wzb-Wzc<sub>R374E</sub>-<br>strep |
| WzcR374E_Rv | GGGTCAGttcGACAATCTCCTGCTGGGTTTTCG |  |
| WzcE354R_Fw | GAAGACcgcAAAGCCAAACTTAACGGTCGC | pET21-Wza-Wzb-Wzc<sub>E354R</sub>-<br>strep |
| WzcE354R_Rv | TGGCTTTgcgGTCTTCCAGCGCCTGACG |  |
| WzcK307E_Fw | GGAAGCAgAAGCGGTGCTCGATTCGATG | pET21-Wza-Wzb-Wzc<sub>K307E</sub>-<br>strep |
| WzcK307E_Rv | CACCGCTTcTGCTTCCAGCGGCAGATCA |  |
| WzcD301R_Fw | TCTGTTcgcCTGCCGCTGGAAGCAAAAG | pET21-Wza-Wzb-Wzc<sub>D301R</sub>-<br>strep |
| WzcD301R_Rv | GCGGCAGgcgAACAGAATCTTTATCCTGACGGAAGG |  |

| Plasmid name | Description | Source/reference |
| --- | --- | --- |
| pET21-Wza-Wzb-Wzc | Contains entire *E. coli* K12 wzabc operon | This study |
| pET21-Wza-strep-Wzb-Wzc | Production of Wza-strep, Wzb, and Wzc protein | This study |
| pET21-Wza-Wzb-Wzc-strep | Production of Wza-strep, Wzb, and Wzc-strep protein | This study |
| pET21-Wza-Wzb-Wz<sub>CK540M</sub>-strep | Production of Wza-strep, Wzb, and Wz<sub>CK540M</sub>-strep protein | This study |
| pET21-Wza-strep-Wzb-WzCK540M-strep | Production of Wza-strep, Wzb, and WzcK540M-strep protein | This study |
| pET21-Wza-Wzb-Wzc-strep | Production of Wza, Wzb, and Wzc-strep protein | This study |
| pET21-Wza-Wzb-WzcK540M-strep | Production of Wza, Wzb, and WzcK540M-strep protein | This study |
| pET21-Wzb-Wzc-strep | Production of Wzb and Wzc-strep protein | This study |
| pET21-Wzb-WzcK540M-strep | Production of Wzb and WzcK540M-strep protein | This study |
| pKD3 | The cat cassette | 3 |
| pKD46 | the $\lambda$ red recombineering | 3 |
| pET28-Avitag-WzcJR | Production of Avitagged Jellyroll domain of Wzc | This study |
| pET28-Avitag-WzcPeri | Production of Avitagged periplasmic domain of Wzc | This study |
| pACYC-BirA | Used in vivo biotinylation of WzCJR and WzCPeri | Gift from Dr. Joop Van den Heuvel |
| pET21-Wza-Wzb-WzcR374E-strep | Production of Wza, Wzb, and Wzc-strep protein | This study |
| pET21-Wza-Wzb-WzcE354R-strep | Production of Wza, Wzb, and WzcE354R-strep protein | This study |
| pET21-Wza-Wzb-WzcD301R-strep | Production of Wza, Wzb, and WzcD301R-strep protein | This study |
| pET21-Wza-Wzb-WzcK307E-strep | Production of Wza, Wzb, and WzcK307E-strep protein | This study |
| pET21-Wza-Wzb-WzCK332E,Y334A,T335A,H338E-strep | Production of Wza, Wzb, and WzCK332E,Y334A,T335A,H338E-strep protein | This study |
| pET21-Wza-Wzb-WzCΔJR-strep | Production of Wza, Wzb, and WzCΔJR-strep protein | This study |
| pET21-Wza-mScarlet3-Wzb-Wzc-sfGFP | Production of Wza-mScarlet3, Wzb, and Wzc-sfGFP | This study |
| pET21-Wza-mScarlet3-Wzb- | Production of Wza-mScarlet3, Wzb, and WzCK332E,Y334A,T335A,H338E-sfGFP | This study |

| Strain name | Genotype or description | source/reference |
| --- | --- | --- |
| *E. coli* Top10 | F- mcrA $\Delta$(mrr-hsdRMS-mcrBC) ϕ80lacZΔM15 ΔlacX74<br>nupG recA1<br>araD139 $\Delta$(ara-leu)7697 galE15 galk16<br>rpsL(Str<sup>R</sup>) endA1 $\lambda$ | Invitrogen |
| *E. coli* JM109 (DE3) | *endA1*, *recA1*, *gyrA96*, *thi*, *hsdR17* (*r<sub>k</sub><sup>-</sup>*, *m<sub>k</sub><sup>+</sup>*), *relA1*, *supE44*, $\lambda^-$, $\Delta$(*lac-proAB*), [*F'*, *traD36*, *proAB*, *lacI<sup>q</sup>*ZΔM15], IDE3 | Promega |
| *E. coli* JM109(DE3)Δwzabc | *E. coli* JM109 (DE3) Δwzabc::cat | This study |
| *E. coli* BL21 Star (DE3) | F<sup>-</sup>*ompT hsdSB* (*r<sub>B</sub><sup>-</sup>*, *m<sub>B</sub><sup>-</sup>*) *gal dcm rne131* (DE3) | Thermo Fisher Scientific |

137 **Table S3.1 Cryo-EM data collection, refinement and validation statistics**

|  | **Wza_C1**<br>**(EMD-**<br>**53598;**<br>**PDB 9R60)** | **Wza_C8**<br>**(EMD-**<br>**53599;**<br>**PDB 9R61)** | **Wzc<sub>K540M</sub>_C**<br>**1 (EMD-**<br>**53600;**<br>**PDB 9R62)** | **Wzc<sub>K540M</sub>_C**<br>**4 (EMD-**<br>**53601;**<br>**PDB 9R63)** | **Wza-**<br>**Wzc<sub>K540M</sub>_C1**<br>**(Conf 0)**<br>**(EMD-53602;**<br>**PDB 9R64)** | **Wza-**<br>**Wzc<sub>K540M</sub>_C8**<br>**(Conf 0)**<br>**(EMD-53603;**<br>**PDB 9R65)** |
| --- | --- | --- | --- | --- | --- | --- |
| **Data**<br>**collection**<br>**and**<br>**processi**<br>**ng** |  |  |  |  |  |  |
| Magnificat<br>ion | 130k | 130k | 130k | 130k | 130k | 130k |
| Voltage<br>(kV) | 200 | 200 | 200 | 200 | 200 | 200 |
| Electron<br>exposure<br>(e<sup>-</sup>/Å<sup>2</sup>) | 40 | 40 | 40 | 40 | 40 | 40 |
| Defocus<br>range<br>(μm) | -0.6 to -2.0 | -0.6 to -2.0 | -0.6 to -2.0 | -0.6 to -2.0 | -0.6 to -2.0 | -0.6 to -2.0 |
| Pixel size<br>(Å) | 0.91 | 0.91 | 0.91 | 0.91 | 0.91 | 0.91 |
| Symmetry<br>imposed | C1 | C8 | C1 | C4 | C1 | C8 |
| Initial<br>particle<br>images<br>(no.) | 1,581,132 | 1,581,132 | 256,861 | 256,861 | 363,741 | 363,741 |
| Final<br>particle<br>images<br>(no.) | 125,938 | 125,938 | 128,284 | 128,284 | 207,837 | 207,837 |
| Map resolution (Å) | 3.0 | 2.7 | 3.4 | 3.2 | 3.5 | 3.2 |
| FSC threshold | 0.143 | 0.143 | 0.143 | 0.143 | 0.143 | 0.143 |
| **Refinement** |  |  |  |  |  |  |
| Initial model used | AlphaFold3 | AlphaFold3 | AlphaFold3 | AlphaFold3 | AlphaFold3 | AlphaFold3 |
| Model resolution (Å) | 3.1 | 2.8 | 3.6 | 3.3 | 3.6 | 3.3 |
| FSC threshold | 0.5 | 0.5 | 0.5 | 0.5 | 0.5 | 0.5 |
| Map sharpening B factor (Å<sup>2</sup>) | −85.3 | −109.9 | −97.8 | −125.1 | −82.5 | −132.5 |
| Model composition |  |  |  |  |  |  |
| Non-hydrogen atoms | 21,616 | 21,616 | 40,312 | 40,312 | 63,984 | 63,984 |
| Protein residues | 2,784 | 2,784 | 5,168 | 5,168 | 8,208 | 8,208 |
| Ligands | 0 | 0 | 8 ADP-Mg<sup>2+</sup> | 8 ADP-Mg<sup>2+</sup> | 8 ADP-Mg<sup>2+</sup> | 8 ADP-Mg<sup>2+</sup> |
| *B* factors (Å<sup>2</sup>) |  |  |  |  |  |  |
| Protein | 54.43 | 30.49 | 61.37 | 91.68 | 56.29 | 92.81 |
| Ligand | - | - | 33.37 | 45.98 | 118.79 | 141.99 |
| R.m.s. deviations |  |  |  |  |  |  |
| Bond lengths (Å) | 0.005 | 0.005 | 0.004 | 0.005 | 0.004 | 0.004 |
| Bond angles (°) | 1.015 | 1.034 | 0.979 | 1.006 | 0.945 | 0.973 |
| Validation |  |  |  |  |  |  |
| MolProbity score | 1.15 | 1.03 | 1.35 | 1.39 | 1.22 | 1.11 |
| Clash score | 2.41 | 1.81 | 2.94 | 2.71 | 3.48 | 2.21 |
| Poor rotamers (%) | 1.05 | 0.88 |  |  | 0.04 | 0.07 |
| Ramachandran plot |  |  |  |  |  |  |
| Favored (%) | 97.43 | 97.54 | 96.11 | 95.23 | 97.64 | 97.45 |
| Allowed (%) | 2.57 | 2.46 | 3.89 | 4.77 | 2.21 | 2.42 |
| Disallowed (%) | 0.00 | 0.00 | 0.00 | 0.00 | 0.15 | 0.14 |

152 **Table S3.2 Cryo-EM data collection, refinement and validation statistics**

|  | Wza-WzCK540M_<br>_C1 | Wza-WzCK540M_<br>C8 | Wza-WzCK540M_<br>C1 | Wza-WzCK540M_<br>C8 | Wza-WzCK540M_<br>C1 | Wza-WzCK540M_<br>C1 | Wza-WzCK540M_<br>C1 |
| --- | --- | --- | --- | --- | --- | --- | --- |
|  | (Conf I) | (Conf I) | (Conf II) | (Conf II) | (Conf III) | (Conf IV) | (Conf V) |
|  | (EMD-53604;<br>PDB<br>9R66) | (EMD-53605;<br>PDB<br>9R67) | (EMD-53606;<br>PDB<br>9R68) | (EMD-53607;<br>PDB<br>9R69) | (EMD-53608;<br>PDB<br>9R6A) | (EMD-53609;<br>PDB<br>9R6B) | (EMD-53610;<br>PDB<br>9R6C) |
| **Data**<br>**collection**<br>**and**<br>**processi**<br>**ng** |  |  |  |  |  |  |  |
| Magnificat<br>ion | 130k | 130k | 130k | 130k | 130k | 130k | 130k |
| Voltage<br>(kV) | 200 | 200 | 200 | 200 | 200 | 200 | 200 |
| Electron<br>exposure<br>(e<sup>-</sup>/Å<sup>2</sup>) | 40 | 40 | 40 | 40 | 40 | 40 | 40 |
| Defocus<br>range<br>(μm) | -0.6 to -<br>2.0 | -0.6 to -<br>2.0 | -0.6 to -<br>2.0 | -0.6 to -<br>2.0 | -0.6 to -<br>2.0 | -0.6 to -<br>2.0 | -0.6 to -<br>2.0 |
| Pixel size<br>(Å) | 0.91 | 0.91 | 0.91 | 0.91 | 0.91 | 0.91 | 0.91 |
| Symmetry<br>imposed | C1 | C8 | C1 | C8 | C1 | C1 | C1 |
| Initial<br>particle<br>images<br>(no.) | 468,471 | 468,471 | 468,471 | 468,471 | 468,471 | 468,471 | 468,471 |
| Final<br>particle<br>images<br>(no.) | 199,355 | 199,355 | 26,950 | 26,950 | 27,501 | 40,425 | 59,128 |
| Map resolution (Å) | 3.8 | 3.4 | 5.8 | 4.2 | 6.3 | 4.6 | 6.7 |
| FSC threshold | 0.143 | 0.143 | 0.143 | 0.143 | 0.143 | 0.143 | 0.143 |
| **Refinement** |  |  |  |  |  |  |  |
| Initial model used | AlphaFold 3 | AlphaFold 3 | AlphaFold 3 | AlphaFold 3 | AlphaFold 3 | AlphaFold 3 | AlphaFold3 |
| Model resolution (Å) | 4.0 | 3.6 | 7.2 | 4.5 | 7.6 | 8.0 | 9.4 |
| FSC threshold | 0.5 | 0.5 | 0.5 | 0.5 | 0.5 | 0.5 | 0.5 |
| Map sharpening B factor (Å<sup>2</sup>) | −100.4 | −132.5 | −203.9 | −103.0 | −260.5 | −76.4 | −377.8 |
| Model composition |  |  |  |  |  |  |  |
| Non-hydrogen atoms | 63,976 | 63,976 | 64,072 | 64,072 | 64,256 | 62,068 | 86,248 |
| Protein residues | 8,208 | 8,208 | 8,232 | 8,232 | 8,248 | 7,968 | 11,072 |
| Ligands | 8 ADP | 8 ADP | 8 ADP | 8 ADP | 8 ADP | 8 ADP | 8 ADP |
| *B* factors (Å<sup>2</sup>) |  |  |  |  |  |  |  |
| Protein | 88.87 | 79.80 | 274.43 | 202.25 | 147.39 | 80.52 | 74.13 |
| Ligand | 132.20 | 130.52 | 172.32 | 89.91 | 89.91 | 130.52 | 69.83 |
| R.m.s. deviations |  |  |  |  |  |  |  |
| Bond lengths (Å) | 0.006 | 0.004 | 0.004 | 0.003 | 0.015 | 0.013 | 0.023 |
| Bond angles (°) | 1.029 | 0.987 | 0.815 | 0.549 | 1.854 | 1.833 | 2.138 |
| Validation |  |  |  |  |  |  |  |
| MolProbity score | 1.43 | 1.26 | 2.01 | 1.56 | 1.51 | 1.10 | 1.51 |
| Clash score | 5.31 | 3.59 | 16.33 | 5.26 | 0.91 | 0.18 | 1.74 |
| Poor rotamers (%) | 0.06 | 0.10 | 0.00 | 0.00 | 2.92 | 1.81 | 2.33 |
| Ramachandran plot |  |  |  |  |  |  |  |
| Favored (%) | 97.19 | 97.42 | 95.73 | 95.90 | 94.34 | 95.23 | 95.61 |
| Allowed (%) | 2.68 | 2.48 | 3.69 | 3.81 | 5.32 | 4.38 | 3.84 |
| Disallowed (%) | 0.14 | 0.10 | 0.59 | 0.29 | 0.34 | 0.39 | 0.55 |
